# Transcriptional dynamics of bread wheat in response to nitrate and phosphate supply reveal functional divergence of genetic factors involved in nitrate and phosphate signaling

**DOI:** 10.1101/551069

**Authors:** Indeewari Dissanayake, Joel Rodriguez-Medina, Siobhan M. Brady, Miloš Tanurdžić

**Affiliations:** School of Biological Sciences, The University of Queensland, QLD 4072, Australia; Department of Plant Biology and Genome Center, University of California Davis, CA95616, USA

**Keywords:** Nitrate, phosphate, wheat, transcriptomics, gene co-expression networks, *HRS1/HHO*, *TGA*

## Abstract

Nitrate (N) and phosphate (P) levels are sensed by plant cells and signaled via local and systemic signaling pathways to modulate plant growth and development. Understanding the genetic basis of these signaling mechanisms is key to future improvement of nutrient use efficiency. While major progress has been made in understanding N and P signaling pathways and their interaction in the model plant Arabidopsis, understanding of transcriptional responses to N and P in a major monocot crop wheat is lacking. Therefore, we investigated gene expression dynamics of wheat roots in response to N and/or P provision using RNA-Seq. We found that nitrate presence is the major trigger for most of the transcriptional response to occur within 24 h, however, we also identified a large array of synergistic transcriptional responses to concomitant supply of N and P. Through gene co-expression analysis, we identified gene co-expression modules prominent in nitrate signaling and metabolism in wheat. Importantly, we identified likely instances of functional divergence in major N-responsive transcription factors families *HRS1/HHO* and *TGA* of wheat from their rice/Arabidopsis homologues. Our work broadens the understanding of wheat N and P transcriptional responses and aids in prioritizing gene candidates for production of wheat varieties that are efficient in nitrogen usage.

## Introduction

Plant development is a highly plastic process, necessitated by the plants’ need to cope with changes in the environment. Availability of nutrients in the soil, for example, modulates plant form and function by changing the root system architecture or shifting the plant’s metabolism. Nitrogen and phosphorus are plant macronutrients absorbed in the form of nitrates (Crawford and Glass 1998) and orthophosphates (Pi) (Schachtman *et al.* 1998; Marschner 2011) respectively. Many soil types lack in both available nitrate (Schlesinger, 1994; Vance 2001) and phosphate (Beadle 1953; Bieleski 1973). Therefore, nitrogen and phosphorus fertilizers are applied to arable lands in large amounts globally where the excess fertilizer usage contributes to environmental pollution (X. Zhang *et al.* 2015; Tilman *et al.* 2002). Hence, understanding the mechanisms mediating plant responses to nutrient supply as well as to nutrient deficiency is critical for the production of new resilient and productive crops that are efficient in nitrogen and phosphorus usage.

Studies on the dicotyledonous model plant *Arabidopsis thaliana* have been key to unraveling the effects of nitrate and phosphate supply/starvation on plant biology and showed that changes in levels of available nitrate and phosphate trigger signaling pathways that modulate root development (Forde 2014; Chiou and Lin 2011). Transcription factors such as NLP7 (Castaings *et al.* 2009; Marchive *et al.* 2013), PHR1 (Rubio *et al.* 2001, Nilsson *et al.* 2007) and HRS1 (Medici *et al.* 2015) and their downstream targets play crucial roles in nitrate and phosphate signaling. For example, NLP7 regulates genes involved in nitrate signaling and nitrate assimilation upon nitrate resupply (Castaings *et al.* 2009; Marchive *et al.* 2013) while PHR1 regulates phosphate starvation responses (Rubio *et al.* 2001, Nilsson *et al.* 2007). Importantly, these studies have shown that nitrate and phosphate signaling are intricately regulated at the transcriptional, post-transcriptional and post-translational levels (reviewed in Krapp *et al.* 2014; Briat *et al.* 2015). Moreover, plant hormones such as auxin, cytokinin and strigolactone are implicated in nitrate and phosphate signaling pathways (reviewed in Krouk 2017; Kapulnik and Koltai 2016; Guan 2017). For example, auxin plays a major role in the lateral root development in response to local nitrate availability through nitrate-mediated regulation of components in the auxin signaling pathway such as auxin receptor AFB3 (Vidal *et al.* 2010) and auxin response factor ARF8 (Lavenus *et al.* 2013).

Several lines of evidence suggest nitrate and phosphate signaling pathways connect through the ubiquitin E3 ligase *NLA* (*NITRATE LIMITATION ADAPTATION*) and GARP family transcription factor *HRS1* (*HYPERSENSITIVE TO LOW PI-ELICITED PRIMARY ROOT SHORTENING 1*) (Kant *et al.* 2011; Medici *et al.* 2015). For example, Arabidopsis *HRS1/HHO* gene family contains seven members and these genes are orthologous to rice *NIGT1* (*Nitrate-Inducible, GARP-type Transcriptional Repressor 1*) (Sawaki *et al.* 2013). The Arabidopsis *HRS1/HHO* gene family and rice *NIGT1* are MYB-related transcription factors from the GOLDEN2-like subgroup of GARP family transcription factors (H. Liu *et al.* 2009; Sawaki *et al.* 2013). Arabidopsis, *HRS1* and *HHO1* are up-regulated in response to nitrate supply within 6 minutes (Krouk *et al.* 2010), while in rice *NIGT1* is induced within 1 h (Sawaki *et al.* 2013). *HRS1/HHO* gene family members have been identified as transcriptional repressors regulating nitrate starvation responses (Maeda *et al.* 2018; Kiba, *et al*, 2018). *AtHHO2* and *AtHHO3* are transcriptionally induced by nitrate albeit to a lesser extent than *AtHRS1/HHO1* (Maeda *et al.* 2018) while *AtHHO5* and *AtHHO6* were validated as nitrate responsive transcription factors (Varala *et al.* 2018).

While Arabidopsis *HRS1* was first characterized by its mutant root phenotype in response to phosphate starvation (H. Liu *et al.* 2009), later studies reported Arabidopsis *HRS1* and its close paralogue *HHO1* (Krouk *et al.* 2010) as well as rice *NIGT1* (Sawaki *et al.* 2013) as being rapidly and specifically transcriptionally induced by nitrate. This was the first evidence that N and P signaling pathways may, in effect, crosstalk, and recently *AtHRS1* was, indeed, shown to be at the nexus of nitrate and phosphate signaling (Medici *et al.* 2015; Maeda *et al.* 2018; Kiba *et al*., 2018) whereby *HRS1* is regulated transcriptionally by nitrate and post-transcriptionally by phosphate. Therefore, delineating exclusive responses to nitrate or phosphate from the responses to the concomitant supply of nitrate and phosphate will facilitate better understanding on the coordination of plant nitrate and phosphate signaling and homeostasis.

Interestingly, it has been reported that the phenotypic changes in root system architecture in response to nitrate and phosphate are different in Arabidopsis relative to monocot species such as rice and maize (Smith and I. De Smet 2012; Niu *et al.* 2013; Shahzad and Amtmann 2017). These phenotypically different responses could be due to underlying gene regulatory mechanisms. Indeed, compelling evidence shows orthologues of a gene performing opposite functions in Arabidopsis and rice. For example, SPX-Major Facility Superfamily3 proteins are implicated in vacuolar transport of phosphate and behave as phosphate influx transporters in Arabidopsis (J. Liu *et al.* 2015) while they are phosphate efflux transporters in rice (C. Wang *et al.* 2015).

Wheat is a major cereal crop accounting for an annual production of 749 million metric tons in 2016 (Food and Agriculture Organization statistics), being second in production to maize (1 billion metric tons in 2016). The hexaploid genetic structure of bread wheat renders it an interesting, yet complex genetic system to study due to its two recent duplication events (Marcussen *et al.* 2014). Evidence for independent regulation of three subgenomes in hexaploid wheat (Pfeifer *et al.* 2014; Powell *et al.* 2017) have been reported. Here, we aim to determine the effects of nitrate and/or phosphate supply on transcriptional dynamics in wheat. Our results show transcriptional reprogramming by nitrate and phosphate provision and delineate the transcriptional responses to nitrate provision that are dependent or independent of phosphate. Moreover, we could also identify instances where wheat gene families of transcription factors involved in nitrate and phosphate signaling such as *TGA* (*TGACG SEQUENCE-SPECIFIC BINDING PROTEIN*) and *HRS1/HHO* (*HYPERSENSITIVE TO LOW PI-ELICITED PRIMARY ROOT SHORTENING 1/HRS1-HOMOLOG*) have functionally diverged from their rice or Arabidopsis homologues. Altogether, our results may contribute to prioritizing gene candidates for future wheat nutrient use efficiency breeding efforts.

## Results and discussion

### Nitrate and/or phosphate provision affects wheat root system architecture

To investigate transcriptome dynamics in response to N and P supply, we first determined if growth under N and/or P deplete or replete conditions had any effect on wheat root system architecture. To this end, we measured root system parameters and *in planta* N and P content. To maximize root responses to N and/or P supply, we grew *Triticum aestivum* (cultivar Chinese spring) plants in water for 11 days. This led to a two-fold and four-fold decrease in nitrogen and phosphorus content in the roots, respectively, relative to plants grown in full strength nutrient solution (see materials and methods) (Figure S1). We then supplied the plants with nutrient solutions which differed in nitrate and phosphate concentrations (0.5 mM or 0 mM phosphate, and/or 2 mM or 0 mM nitrate). We therefore used four treatment combinations (-P-N, −P+N, +P-N and +P+N), here denoted as pn, pN, Pn and PN, respectively. After further growing the plants under these four conditions for seven days we were able to detect changes in root system architecture (Figure 1A).

**Figure 1.**
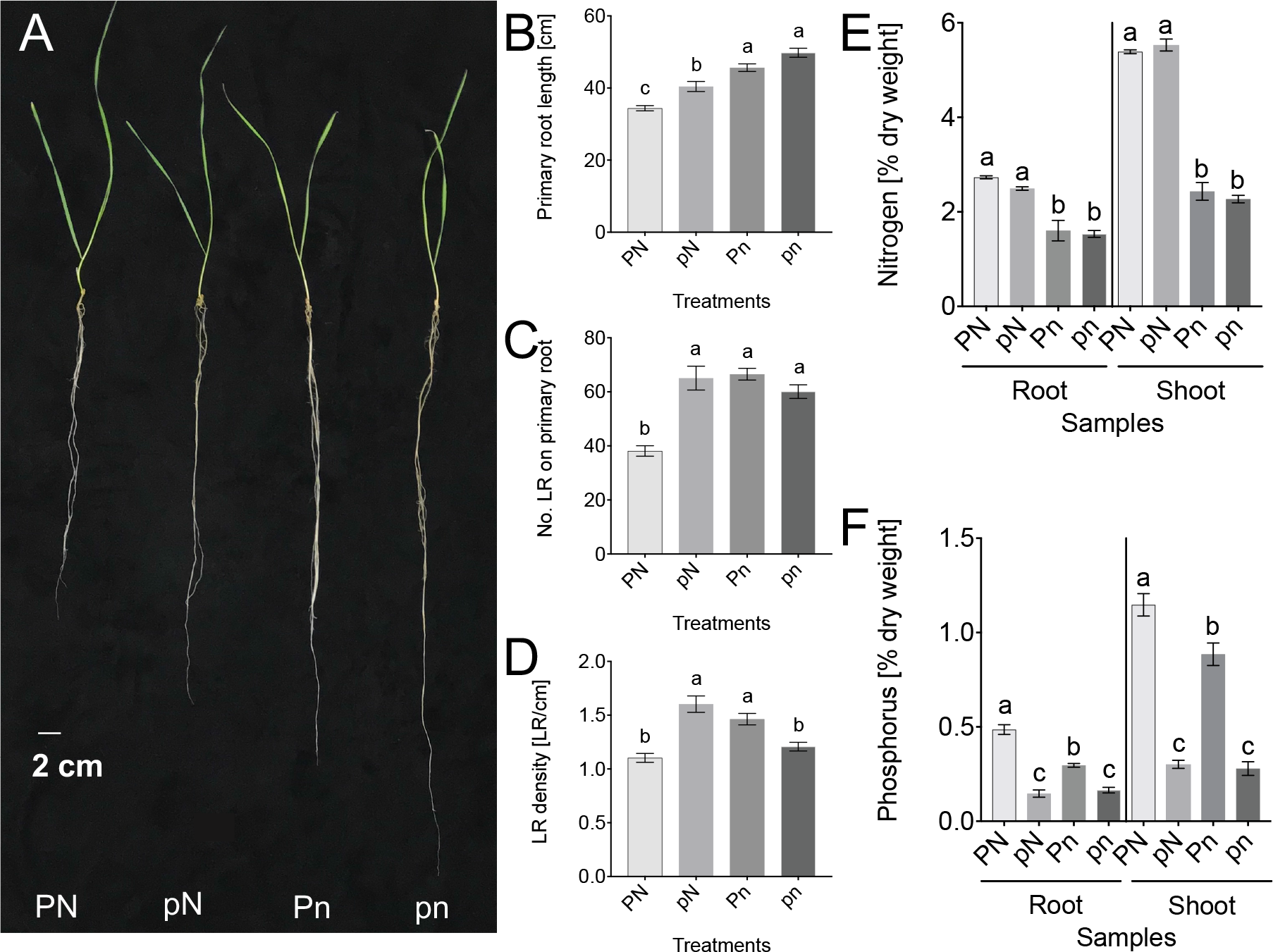
The wheat root responses to nitrate and/or phosphate supply. **(A)** Root system architecture of wheat plants treated with nitrate and/or phosphate. Comparison of wheat root system parameters **(B)** primary root length, **(C)** number of lateral roots (LR) on the primary root, **(D)** lateral root (LR) density on the primary root (n=15, error bars indicate standard error of mean). Comparison of **(E)** Nitrogen content, **(F)** Phosphorus content of total root and total shoot tissue (n=3, error bars indicate standard error of mean). Figures 1A-F are results from 18-day old wheat plants that were grown in indicated nutrient regimes for seven days, letters denote significance based on two-way ANOVA followed by Tukey HSD (*P*<0.05) in Figures 1B-F.

Following N and/or P supply for 7 days, we measured N and P content in roots and shoots of 18 days old plants (Figure 1E-F). As expected, N content was significantly higher in roots and shoots of plants grown in pN and PN conditions than in Pn and pn (Figure 1E). Phosphorus content was significantly higher in roots and shoots of plants grown in Pn and PN conditions, than in the plants grown in pN and pn conditions. Interestingly, the phosphorus content was also significantly lower in Pn treated plants than in PN treated plants (Figure 1F). This result first suggested that presence of nitrogen may facilitate phosphorus uptake/redistribution because an equal amount of phosphate was supplied in Pn and PN treatments.

Plant root system architecture is modulated by nutrient resources available through nutrient sensing and signaling (Kellermeier *et al.* 2014; Shahzad and Amtmann 2017). We measured three root system parameters (primary root length, the number of lateral roots and lateral root density on the primary root) of 18 day old plants, as described above. The three root system parameters were significantly different among treatments (Figure 1A-D): single (pN, Pn) or combined (pn) nutrient deficiency led to increased primary root length (Figure 1B) and increased number of lateral roots on the primary root (Figure 1C), relative to the nutrient replete (PN) condition. Single nutrient deficiency also led to increased lateral root density, suggestive of root foraging for nitrate and phosphate. However, both PN and pn treatments resulted in similar lateral root densities (Figure 1D). These results showed that wheat roots under single P/N deficient conditions employ strategies for increasing root mass by both increasing main root length and the number of lateral roots. They also suggest that N availability supersedes the effect of P availability.

Previous studies of Arabidopsis root responses to phosphate deficiency reported shorter primary roots and increased lateral root density. This is thought to be a mechanism for efficient top-soil foraging (Péret *et al.* 2014), although the responses may vary depending on the severity and the exposure time to the phosphate deficiency (Niu *et al.* 2013; Tian *et al.* 2014) and the genetic background since some natural accessions of *Arabidopsis thaliana* show difference in P response (Chevalier *et al.* 2003; Shahzad *et al.* 2018). Similarly, root responses to nitrate deficiency vary depending on the severity of deficiency as well as the distribution of nitrate (homogeneous vs local distribution in soil) (Gruber *et al.* 2013, reviewed in Giehl *et al.* 2013). Under severe nitrate deficiency conditions, both primary and lateral root growth were retarded in Arabidopsis (Giehl *et al.* 2013; Gruber *et al.* 2013; Giehl and Wirén 2014). Studies on rice reported increased seminal root length and decreased lateral root density (Sun *et al.* 2014) due to nitrate or phosphate deficiency. Our results on wheat primary root length under single nutrient deficiency are in agreement with those reported in the study on rice (Sun *et al.* 2014; Yu *et al.* 2016), however they differ in lateral root density where nitrate or phosphate deficiency caused increased lateral root density in wheat. This could be due to the differences in the nitrogen and phosphorus concentrations used in our experiment and the study by Sun *et al.* 2014.

### Analysis of differential gene expression reveals major combinatorial effects of nitrate and phosphate provision on transcriptional dynamics

Since rapid transcriptomic changes have been observed in response to nitrate (Krouk *et al.* 2010; K.-H. Liu *et al.* 2017), and phosphate supply (Gutiérrez-Alanís *et al.* 2017; Secco *et al.* 2013), we focused on exploring the dynamics of wheat root transcriptional responses within the first 24 h of nitrate and/or phosphate supply. Moreover, considering the emerging evidence of a connection between nitrate and phosphate signaling pathways (Maeda *et al.* 2018; Kiba *et al*., 2018) as well as our results on root system architecture changes in wheat roots in response to combinatorial nitrate and phosphate treatment, we used an experimental setup that enabled delineating combinatorial effects of nitrate and phosphate on the root transcriptome from the effects due to nitrate- or phosphate-only.

To characterize transcriptomic responses to P and N provision, we collected root samples at 1 h, 2 h, 4 h and 24 h following nutrient supply and quantified gene expression using RNA-Seq. Out of 110790 wheat high confidence genes in IWGSC RefSeq version 1.0 annotation, 94253 (85.03%) had non-zero total read count across samples. We herein denote these 94253 genes as “expressed” under at least one of the conditions.

We considered the numbers of differentially expressed genes (FDR < 0.05) between a treatment (PN or Pn or pN) and reference (pn) at four time points (Figure 2A). Only 25 genes (0.02% of expressed genes) responded to phosphate supply in the absence of nitrate (Pn) within the first 24 h (Table S8). These 25 genes included the SPX domain containing gene most similar to *SPX DOMAIN GENE 3* in Arabidopsis (*TraesCS2D01G177100*, orthologous to *ATSPX3* (*AT2G45130*)) down-regulated at 1 h; three phosphate transporter genes (*TraesCS4B01G317200*, *TraesCS4A01G416500* and *TraesCSU01G070800* (orthologous to *ATPT1* (*PHOSPHATE TRANSPORTER 1*, *AT5G43350*)) down-regulated at 2 h, and *CLE* gene family members (*TraesCS1D01G401200* and *TraesCS1B01G421600*, up-regulated at 24 h). Transcriptional repression of both *ATSPX3* (Duan *et al.* 2008) and *ATPT1* (Puga *et al.* 2014) by PHR1 under P sufficient conditions has been reported in Arabidopsis (reviewed in Gu *et al.* 2016). This result suggests that despite few wheat genes are regulated by phosphate in the absence of nitrate, at least some components of the PHR1 signaling module are conserved in function between wheat and Arabidopsis and respond rapidly to phosphate supply independent of nitrate. Even though *IPS1* and *miR399* are also well known targets of PHR1, these genes could not be assessed using RNA-Seq since these ncRNA loci are not present in the current wheat high confidence gene annotation.

**Figure 2.**
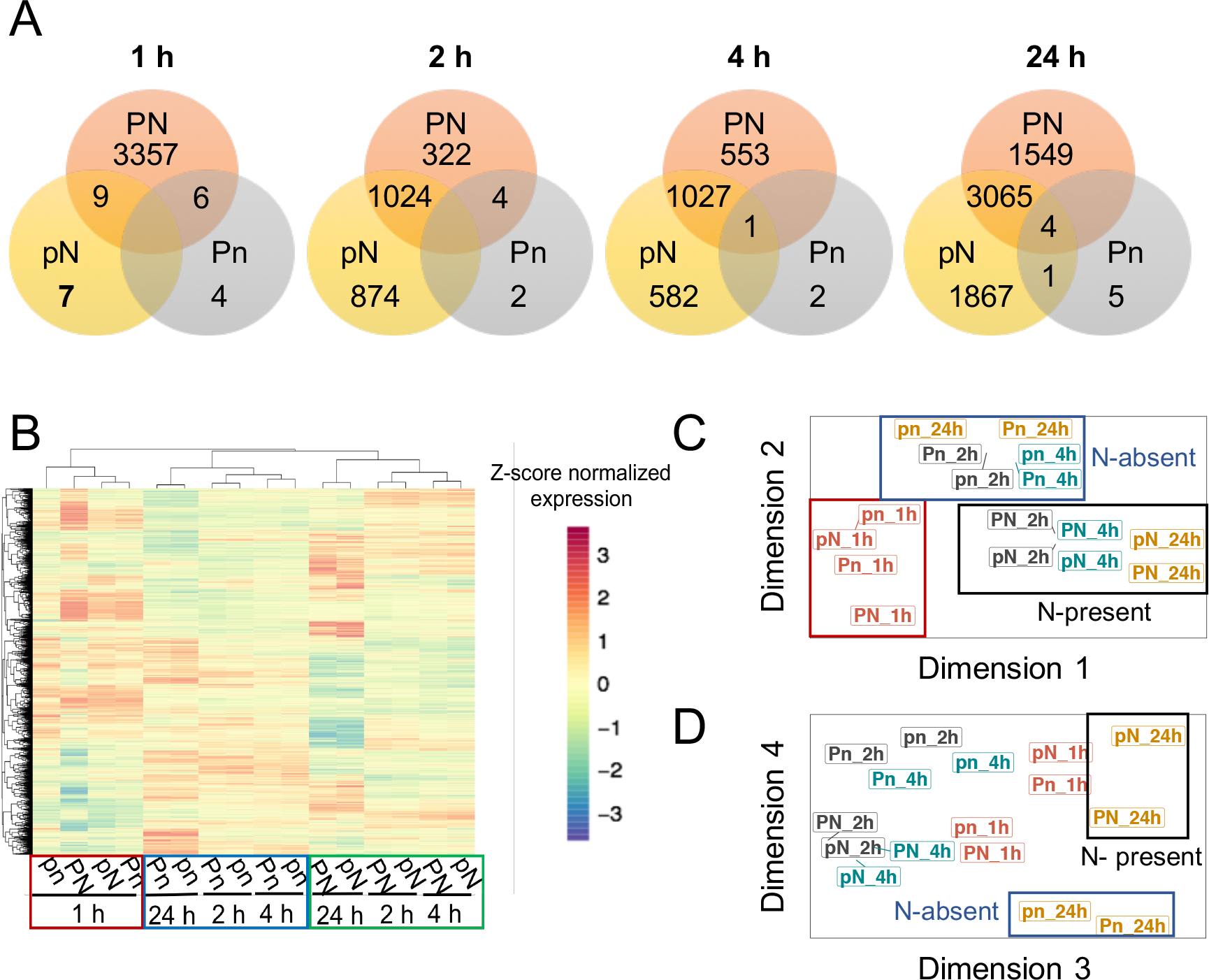
Combinatorial effects of nitrate and phosphate provision in modulating gene expression. **(A)** Venn diagrams showing the overlap of differentially expressed (DE) genes among different treatment comparisons with respect to pn treatment within a time point. **(B)** Hierarchical clustering of the DE genes based on z-score normalized expression values. Multi-dimensional scaling plots representing variance as vectors in the first four dimensions **(C)** dimensions 1 and 2; **(D)** dimensions 3 and 4.

Following N supply in the absence of P (pN treatment) for 1 h, 16 genes (0.01% of expressed genes) were differentially expressed (Figure 2A, Table S8), only two of which were up-regulated (Table S2). This changed dramatically at later time points with thousands of genes being differentially expressed under pN treatment (Figure 2A, Table S2), starting with the the 2 h time point. These results show the progression of a transcriptional cascade related to nitrate signaling and metabolism (further discussed below).

Strikingly, when nitrate and phosphate were supplied concomitantly (PN), massive transcriptome changes occur as early as 1 h, with 3372 genes (3.5% of expressed genes) differentially expressed (1514 up-regulated, 1858 down-regulated) (Figure 2A, Table S2). This transcriptional response subsides at 2 h and 4 h with 1350 (1.4% of expressed genes) and 1581 (1.6% of expressed genes) genes differentially expressed, respectively. Another wave of transcriptional response could then be detected at 24 h, with 4618 genes (4.8% of expressed genes) differentially expressed (2080 up-regulated and 2538 down-regulated) in response to PN treatment (Figure 2A, Table S2, Table S8). Altogether, by considering gene expression across the three treatments (pN, Pn and PN relative to pn) and the four time points, 10229 genes (10.8% of expressed genes) (Table S8) were identified as differentially expressed at least in one treatment-time point combination.

To identify sub-groups within 10229 differentially expressed genes based on the similarity in expression profiles in an unsupervised manner, we carried out a hierarchical clustering analysis (Figure 2B). Intriguingly, Pearson correlation-based clustering of variance-stabilized read counts showed that treatments at 1 h induce gene expression changes forming a separate cluster from rest of the treatment-time combinations. This observation implies that early responses to P and N are distinct from the transcriptomic responses at later time points. Furthermore, if P and N signaling pathways were mutually independent, we would expect to see the clustering based on treatment, regardless of the time points. However, the clustering pattern suggests that wheat root responses to P supply are dependent on the supply of N. Our results indicate the combinatorial effects of nitrate and phosphate provision in modulating gene expression and validate our approach that took into account the interaction effects in response to P and N supply.

To identify the contribution from time factor and treatment to the variation in expression profiles we analyzed multi-dimensional scaling (MDS) plots (Figure 2C, 2D, Figure S2A). The first four dimensions explain 85.7% of the variance (Figure S2A) where first two dimensions essentially recapitulate the patterns observed with hierarchical clustering (Figure 2C). While dimension 3 accounts for variance due to time factor, dimension 4 explains variance due to presence or absence of nitrate (Figure 2D). Considering both the overlap of differentially expressed genes (Figure 2A) and the clustering pattern of the treatment comparisons (Figure 2B-2D) we conclude that while concomitant supply of nitrate and phosphate have a synergistic effect on the transcriptomic dynamics occurring at 1 h, at later time points, the presence of nitrate is required for most of the transcriptional response to occur.

While the bulk of gene expression changes that require supply of both P and N (PN treatment) happen by 1 h (Figure 2A), the number of genes that respond to N supply regardless of P status (the overlap between pN and PN treatments) gradually increases over time from 9 to 3069 genes (Figure 2A). If a gene is differentially expressed in more than one treatment, directionality of fold change (up- or down-regulation) could indicate whether the gene is transcriptionally regulated the same way by the different treatments. To this end, directionality of fold change (up/down regulation with respect to pn condition) remained the same in all the genes that were differentially expressed in both pN and PN conditions at a given time point (the overlap between pN and PN treatments), except for *TraesCS6A01G165100*, which was transcriptionally upregulated by N (PN and pN), however the supply of P in the absence of N (Pn) led to its down-regulation at 4 h after treatment (Table S8). This gene is annotated as a nicotianamine synthase in wheat. Congruent with our observations, previous studies have reported up-regulation of nicotianamine synthase genes in response to nitrate supply in Arabidopsis (R. Wang *et al.* 2000; R. Wang *et al.* 2003). However, in the study by (Shukla *et al.* 2017), they reported up-regulation of *NICOTIANAMINE SYNTHASE 2* in response to excess phosphate levels (20 mM phosphate and 3.9 mM nitrate) in Arabidopsis. Our results show that wheat nicotianamine synthase *TraesCS6A01G165100* is regulated similarly only in the presence of N. This suggests that *TraesCS6A01G165100* is regulated by phosphate in a nitrate dependent manner. Since nicotianamine synthase is involved in iron transport (Schuler *et al.* 2012), our results suggest nitrate and phosphate-dependent effects on iron distribution through nicotianamine synthase.

The majority of the studies characterizing nitrate-induced transcriptional responses in Arabidopsis have been carried out in the presence of sufficient phosphate. Considering the emerging evidence of crosstalk between nitrate and phosphate signaling (Kant *et al.* 2011; Medici *et al.* 2015, Maeda *et al.* 2018), our approach enables delineation of the responses to nitrate that are dependent or independent of presence of phosphate. To that end, we performed GO term enrichment analysis using genes that were differentially expressed exclusively in pN vs pn or in PN vs pn treatments at each of the four time points (Table 1, Table S3).

**Table 1.** Lowest level GO terms enriched in pN vs pn and PN vs pn treatment comparisons, along with the fold enrichment (All GO terms shown were enriched with FDR *P*<0.05).

| Treatment-time combination | GO category | Fold enrichment |
| --- | --- | --- |
| pN_1 h down | phosphatidylcholine biosynthetic process | 1151.16 |
| pN_2 h up | glutamate biosynthetic process | 138.48 |
|  | nitrate transport | 69.24 |
|  | histidine biosynthetic process | 39.57 |
|  | NAD biosynthetic process | 27.70 |
|  | chloride transport | 24.44 |
|  | cytokinin metabolic process | 15.39 |
|  | zinc ion transmembrane transport | 14.58 |
|  | regulation of cyclin-dependent protein kinase activity | 14.20 |
|  | response to auxin stimulus | 4.48 |
| pN_2 h down | L-phenylalanine catabolic process | 30.15 |
|  | fatty acid beta-oxidation | 29.61 |
|  | glutathione catabolic process | 23.69 |
|  | amino acid transmembrane transport | 7.05 |
|  | response to wounding | 4.51 |
|  | oxidation reduction | 1.59 |
| pN_4 h up | maturation of SSU-rRNA from tricistronic rRNA transcript (SSU-rRNA, 5.8S rRNA, LSU-rRNA) | 121.57 |
|  | intracellular distribution of mitochondria | 91.18 |
|  | glutamate biosynthetic process | 60.79 |
|  | negative regulation of ethylene mediated signaling pathway | 45.59 |
|  | isoleucine biosynthetic process | 40.52 |
|  | proline biosynthetic process | 39.08 |
|  | glutathione catabolic process | 24.31 |
|  | histidine biosynthetic process | 17.37 |
|  | defense response to fungus | 17.37 |
|  | response to desiccation | 15.20 |
|  | arginine biosynthetic process | 14.59 |
|  | tricarboxylic acid cycle | 14.25 |
|  | glycolysis | 10.80 |
|  | trehalose metabolic process | 9.27 |
|  | protein amino acid dephosphorylation | 8.82 |
|  | oligosaccharide biosynthetic process | 8.05 |
|  | sulfur amino acid biosynthetic process | 6.59 |
| pN_4 h down | mitochondrial pyruvate transport | 74.42 |
|  | cell proliferation | 62.02 |
|  | heparan sulfate proteoglycan biosynthetic process | 62.02 |
|  | glycosaminoglycan biosynthetic process | 53.16 |
|  | response to wounding | 20.52 |
| pN_24 h up | photosystem II stabilization | 34.20 |
|  | pyrimidine nucleotide biosynthetic process | 25.65 |
|  | glycolipid biosynthetic process | 25.65 |
|  | response to nitrate | 25.65 |
|  | cellular iron ion homeostasis | 20.52 |
|  | nitrate transport | 20.52 |
|  | phosphatidylcholine biosynthetic process | 11.40 |
|  | galactose metabolic process | 8.02 |
| pN_24 h down | S-adenosylmethionine biosynthetic process | 27.18 |
|  | L-serine biosynthetic process | 16.56 |
|  | L-phenylalanine catabolic process | 14.46 |
|  | galactose metabolic process | 13.09 |
|  | cell growth | 11.62 |
|  | cellulose microfibril organization | 11.62 |
|  | aromatic amino acid family biosynthetic process | 11.36 |
|  | activation of protein kinase C activity by G-protein coupled receptor protein signaling pathway | 11.04 |
|  | cellulose biosynthetic process | 8.76 |
|  | zinc ion transmembrane transport | 6.97 |
|  | chitin catabolic process | 5.74 |
|  | nicotinamide nucleotide metabolic process | 5.27 |
|  | hemicellulose metabolic process | 3.51 |
|  | cellular cell wall macromolecule metabolic process | 3.46 |
| PN_1 h up | protein sumoylation | 55.90 |
|  | glutamate biosynthetic process | 55.90 |
|  | intracellular distribution of mitochondria | 55.90 |
|  | ribosomal large subunit assembly | 37.27 |
|  | phosphatidylserine biosynthetic process | 27.95 |
|  | nitrate transport | 20.96 |
|  | glutamyl-tRNA aminoacylation | 15.24 |
|  | gluconeogenesis | 13.97 |
|  | threonyl-tRNA aminoacylation | 12.42 |
|  | pentose-phosphate shunt | 9.58 |
|  | malate transport | 9.32 |
|  | NAD biosynthetic process | 8.38 |
|  | chloride transport | 8.22 |
|  | histidine biosynthetic process | 7.99 |
|  | protein deubiquitination | 6.82 |
|  | fructose 6-phosphate metabolic process | 6.78 |
|  | glycolysis | 6.25 |
|  | calcium ion transmembrane transport | 6.21 |
|  | trehalose biosynthetic process | 5.99 |
|  | sodium ion transport | 5.59 |
|  | ubiquitin-dependent protein catabolic process | 2.01 |
|  | RNA processing | 1.93 |
| PN_1 h down | L-serine biosynthetic process | 8.77 |
|  | glucosylceramide catabolic process | 8.77 |
|  | malate transport | 7.79 |
|  | regulation of cyclin-dependent protein kinase activity | 5.99 |
|  | L-phenylalanine catabolic process | 5.10 |
|  | oligopeptide transport | 4.73 |
|  | xyloglucan metabolic process | 4.11 |
|  | plant-type cell wall organization | 3.51 |
|  | response to oxidative stress | 3.36 |
| PN_2 h up | malate transport | 47.96 |
|  | zinc ion transmembrane transport | 30.29 |
|  | regulation of response to stimulus | 25.03 |
|  | plant-type cell wall organization | 14.39 |
| PN_2 h down | transmembrane transport | 3.07 |
| PN_4 h up | rRNA methylation | 98.23 |
|  | nitrogen compound transport | 42.10 |
|  | pseudouridine synthesis | 12.63 |
|  | metal ion transport | 5.59 |
|  | ammonium transport | 28.07 |
| PN_4 h down | abscisic acid biosynthetic process | 87.09 |
|  | allantoin catabolic process | 87.09 |
|  | phosphate transport | 38.42 |
|  | valyl-tRNA aminoacylation | 34.83 |
|  | sulfate transport | 15.83 |
|  | L-phenylalanine catabolic process | 14.25 |
|  | respiratory electron transport chain | 9.11 |
|  | ATP synthesis coupled proton transport | 7.46 |
|  | recognition of pollen | 3.61 |
| PN_24 h up | glutamate biosynthetic process | 71.39 |
|  | threonine biosynthetic process | 42.83 |
|  | acetyl-CoA biosynthetic process from pyruvate | 21.42 |
|  | phosphatidylcholine biosynthetic process | 21.42 |
|  | isoleucine biosynthetic process | 19.04 |
|  | L-methionine salvage from methylthioadenosine | 17.13 |
|  | trehalose biosynthetic process | 13.77 |
|  | nicotianamine biosynthetic process | 12.24 |
|  | malate transport | 10.71 |
|  | tricarboxylic acid cycle | 10.71 |
|  | cytokinin metabolic process | 9.52 |
|  | sulfate transport | 7.79 |
|  | glycolysis | 7.33 |
|  | nucleoside metabolic process | 5.91 |
|  | lipid transport | 4.56 |
| PN_24 h down | cellular iron ion homeostasis | 35.08 |
|  | mitochondrial pyruvate transport | 21.05 |
|  | oligopeptide transport | 7.10 |
|  | amino acid transmembrane transport | 6.26 |
|  | drug transmembrane transport | 3.76 |
|  | lipid metabolic process | 2.04 |
|  | oxidation reduction | 1.38 |

After 2 h of N supply in the absence of P (pN treatment at 2 h), the over-represented GO terms in the differentially expressed gene set were related to hormonal regulation (“cytokinin metabolism”, “response to auxin stimulus”) (Table 1, Table S3). Specifically, the genes that were up-regulated and assigned to “cytokinin metabolism” GO category at 2 h in response to pN treatment were identified as cytokinin oxidase in wheat (Table S3). This observation is of interest, since cytokinin has been identified as a key hormone in systemic nitrate signaling (Sakakibara *et al.* 2006) in Arabidopsis. Cytokinin is involved in lateral root development in response to systemic nitrate status by acting as a reporter of nitrate demand of the whole plant (Ruffel *et al.* 2011; Ruffel *et al.* 2016). Nitrate is capable of inducing cytokinin oxidases (R. Wang *et al.* 2003; Scheible *et al.* 2004), which in turn regulates cytokinin levels by negative feedback mechanisms (Kieber and Schaller 2018). A recent study suggested a role for cytokinin oxidases in modulating shoot meristem growth triggered by nitrate supply (Landrein *et al.* 2018). Therefore, up-regulation of cytokinin oxidases indicates that by two hours after N supply (in the absence of phosphate) N signaling already is likely to affect cytokinin levels in the roots and, consequently, cytokinin signals moving to the shoot. The genes that were up-regulated in pN treatment and assigned “response to auxin stimulus” GO term are homologous to SAUR-like auxin responsive family genes in Arabidopsis (Table S3). Up-regulation of SAUR-like auxin responsive family members may suggest that by two hours of nitrate supply, auxin levels are also changing in the roots as SAUR gene family members are rapidly induced by auxin (McClure *et al.* 1989; Abel and Theologis 1996). At 4 h, biological processes related to amino acid biosynthesis are enriched in the pN up-regulated gene set (Table 1) and these results are in agreement with previous reports on transcriptomic responses to nitrate in the presence of phosphate (R. Wang *et al.* 2003; Scheible *et al.* 2004). However, our results show that in wheat, these processes occur independent of P status.

Interestingly, cell wall structure related processes such as “cellulose biosynthesis” and “microfibril organization” were enriched in the down-regulated gene sets in response to pN, specially at 24 h (Table1, Table S3). This may suggest that upon nitrate supply, cell division and expansion which ensures longer roots in the search for nitrate is reduced. Five genes (*TraesCS5D01G401900, TraesCS5A01G095200, TraesCS5A01G392000, TraesCS5B01G396900* and *TraesCS5D01G107900*) specifically down-regulated in response to nitrate supply at 24 h were assigned GO terms of “cellulose microfibril organization” and “cell growth” (Table S3) and have been annotated as COBRA-like proteins in wheat. *COBRA* is a plant-specific glycosylphosphatidylinositol-anchored protein and is upregulated in cells entering the zone of elongation (Roudier 2002) in Arabidopsis. The involvement of *COBRA* gene family members in shaping up plant root system architecture has been studied in Arabidopsis (Brady *et al.* 2006; Schindelman *et al*., 2001; Benfey and Scheres 2000). The five COBRA-like wheat genes that are specifically down-regulated at 24 h of nitrate supply are orthologous to *COBRA* and *COBRA-Like 4* in Arabidopsis and *BRITTLE STALK-2* (*Bk2*), *BRITTLE STALK-LIKE*-*3* (*Bk2L3*) and *BRITTLE STALK-LIKE*-*4* (*Bk2L4*) in maize (Figure S2B). *Bk2L4* is down-regulated within 2 h of nitrate starvation, while *Bk2L3* remained unresponsive to nitrate starvation (Brady *et al.* 2006). While both wheat and maize *COBRA* gene orthologues respond to nitrate (supply or starvation), it is intriguing that the down-regulation of wheat *Bk2L3/4* orthologues in response to nitrate supply happens only in the absence of phosphate at 24 h.

The biological processes enriched in the up-regulated gene category at 1 h in response to PN include carbon metabolism and transport of ions such as nitrate and calcium (Table 1, Table S3). Some of the processes, such as “RNA processing” and “pentose phosphate shunt”, were previously reported to be enriched among the genes up-regulated in response to nitrate supply in the presence of phosphate (R. Wang *et al.* 2003; Scheible *et al.* 2004; Krouk *et al.* 2010). Interestingly all the genes that were assigned GO term “malate transport” were annotated as aluminum activated malate transporter family members in wheat (Table S3). Some of these genes (*TraesCS7B01G127200, TraesCS7B01G116000, TraesCS7A01G208700, TraesCS7D01G211100*) were up-regulated while some (*TraesCS6B01G270300, TraesCS6D01G236500, TraesCS2D01G399700*, *TraesCS2B01G420600*) were down-regulated as early as 1 h (Table 1, Table S3). Later in the time course, aluminum activated malate transporter genes were found up-regulated by PN treatment at 2 h and 24 h. Interestingly, wheat aluminum activated malate transporters have been found to be permeable not only to malate, but also to nitrate and chloride (Piñeros *et al.* 2008; W.-H. Zhang *et al.* 2008). Moreover, AtALMT1 (ALUMINUM ACTIVATED MALATE TRANSPORTER 1) is a key regulator in modulating root development under phosphate deficiency (Mora-Macías *et al.* 2017; Balzergue *et al.* 2017). Therefore, our results suggest involvement of aluminum activated malate transporters in response to nitrate and phosphate, although their precise function (as nitrate transporter or functioning in P starvation induced root development) remains unresolved.

Moreover, wheat genes annotated as ubiquitin carboxyl-terminal hydrolases were enriched in GO term categories “ubiquitin dependent protein catabolic process” and “protein deubiquitination” (Table 1, Table S3). The Arabidopsis homologues of these genes are members of ubiquitin specific protease (UBP) family (Table S3). While UBP14 is involved in the adaptation of root development to local phosphate availability (W.-F. Li *et al.* 2010), none of the Arabidopsis homologs in these GO categories were UBP14 (Table S3). Interestingly, nicotianamine synthase genes and Yellow Stripe like (YSL) genes were present in the over-represented GO categories “nicotianamine biosynthetic process” and “transmembrane transport”. Considering nicotianamine synthase (Koen *et al.* 2013) and YSL genes (Waters *et al.* 2006; Kumar *et al.* 2017) are involved in plant iron homeostasis, these results may suggest the changes in the iron homeostasis are affected by concomitant supply of N and P. Moreover, wheat phosphate transporters which are homologous to *AtPT1* and *AtPT1;4* were down-regulated at 4 h, suggesting that plants experience phosphate sufficient conditions by this time, thereby repressing the phosphate transporters. While four cytokinin oxidase genes were present in the “cytokinin metabolic process” category for genes up-regulated at 24 h (Table S3), these genes were not present in the cytokinin oxidase gene set that were responding to pN, suggesting the activity of some wheat cytokinin oxidases may depend on the presence of phosphate.

### Gene co-expression network analysis reveals gene modules involved in nitrate signaling in wheat

Correlation based methods using similarity of gene expression profiles to infer functions of the genes through guilt-by-association principle have been widely employed in systems biology research (Wolfe *et al.* 2005; Lee *et al.* 2010; Usadel *et al.* 2009; R. De Smet and Marchal 2010; Banf and Rhee 2017). To perform gene co-expression analysis using WGCNA (Langfelder and Horvath 2008), we selected 8300 genes with the highest variance in gene expression across the entire dataset (top 10%). The gene co-expression module-treatment (“module-trait”) relationships that resulted from weighted gene co-expression analysis are shown in Figure 3A and were used to identify modules for further analyses.

**Figure 3.**
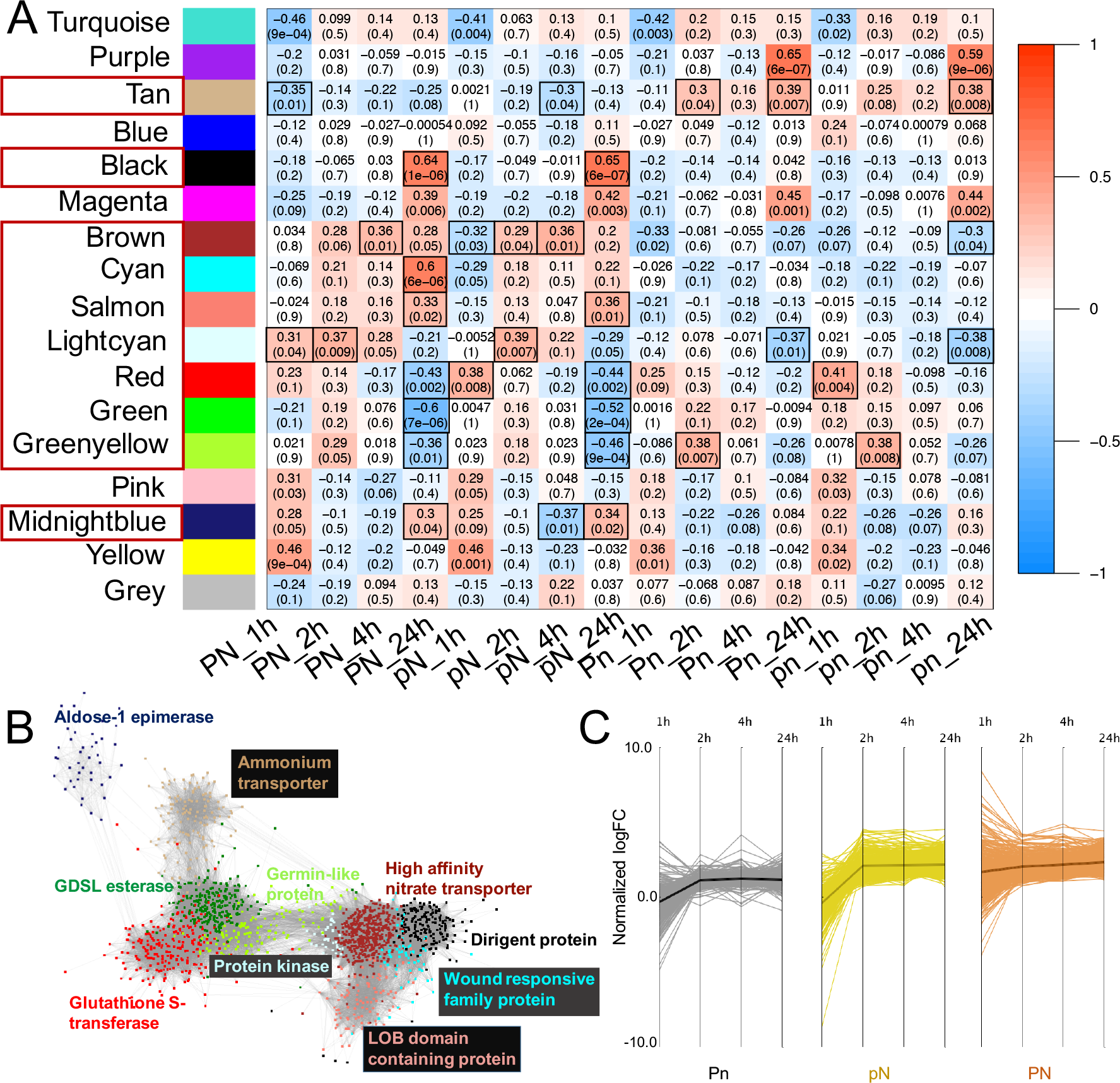
Weighted gene co-expression network (WGCNA) analysis identified gene modules involved in nitrate signaling in wheat. **(A)** Heatmap showing expression between network modules and the treatments (traits). Module eigengenes were calculated using WGCNA package and each cell shows the corresponding correlation value and its *P*-value. Color scale corresponds to the correlation values. Cells outlined in black represent the module-trait relationships selected for further analysis. **(B)** The hub genes were identified based on intramodular connectivity. **(C)** Gene expression profile for the Brown module; Y axis represents the normalized log fold change (logFC) values where expression was measured as treatment/reference (i.e. pn) for a given time point; mean (black or gold or orange line) is shown for all the genes (shaded lines) in the module.

To delineate the independent effects of nitrate or phosphate on gene co-expression from those due to combined nutrients, we selected the modules that showed significant correlation coefficient values in a treatment-dependent pattern. This resulted in a sub-network with ten modules (tan, lightcyan, brown, midnightblue, black, cyan, salmon, red, green and greenyellow modules) (Figure 3A). We identified the top 5 hub genes in each module based on intramodular connectivity. In three of the modules in the sub-network, homeologues of a gene were among the hub genes where black module contained homeologue triad of a dirigent protein, salmon module contained a homeologue triad of a putative LOB domain containing protein and brown module contained homeologue duplet of a high affinity nitrate transporter (Table S4). The functional annotation for these hub genes identified based on intramodular connectivity are shown in Figure 3B. Moreover, to identify the biological processes enriched in each module we carried out a GO term enrichment analysis on the modules. The top 5 GO terms enriched in each module are listed in Table S4. According to this analysis, the brown module contains genes with a function related to nitrate signaling and/or transport (Figure 3B, Table S4). The genes in the brown module also showed expression profiles with higher expression under pN and PN treatments throughout the time course (Figure 3C).

Based on the principle of guilt-by-association, we predicted the transcription factors co-expressed with the genes in the brown module could play a regulatory role in nitrate signaling/metabolism. According to the annotation available for the wheat genome (International Wheat Genome Sequencing Consortium (IWGSC) *et al.* 2018), the brown module contained 23 transcription factors/transcription factor-like proteins which included homeologue triads of MYB transcription factors (*TraesCS2A01G488200*, *TraesCS2B01G515800*, *TraesCS2D01G488500* and *TraesCS5A01G401600*, *TraesCS5B01G406300*, *TraesCS5D01G411800*), transcription factor-like proteins (*TraesCS1A01G276600*, *TraesCS1B01G285800*, *TraesCS1D01G276100*) (Table S5). Out of these 23 transcription factors, the two homeologue triads *TraesCS5A01G401600*, *TraesCS5B01G406300*, *TraesCS5D01G411800* and *TraesCS1A01G276600*, *TraesCS1B01G285800*, *TraesCS1D01G276100* have Arabidopsis orthologues that are members of *HRS1/HHO* and *TGA* gene families, respectively. Since some *HRS1/HHO* ici *et al.* 2015) and *TGA* (Álvarez *et al.* 2014) family members are involved in nitrate ling and metabolism, we decided to further probe the role of these two homeologue in the context of nitrate signaling in wheat.

### Wheat *HRS1/HHO* and *TGA* family members show functional divergence from Arabidopsis orthologues in response to nitrate

Given the role of the *HRS1/HHO* gene family in response to N and P in Arabidopsis and rice (Medici *et al.* 2015; Maeda *et al.* 2018; Kiba *et al.*, 2018), we explored gene expression dynamics of the wheat *HRS1/HHO* gene family in response to nitrate and phosphate supply. Firstly, we identified putative wheat *HRS1/HHO* orthologues by protein sequence alignment using reciprocal BLAST. This confirmed that all the wheat *HRS1/HHO* sequences contain the conserved G2-like DNA binding domain (Figure S3A). A maximum likelihood-built gene tree showed that a single wheat homeologue triad falls within the *AtHRS1* and *OsNIGT1* clade, another homeologue triad groups within the *AtHHO4* clade, while three homeologue triads cluster within the *AtHHO5/HHO6* clade (Figure 4A). Of those 15 wheat *HRS1/HHO* orthologues, only 10 genes were found to be differentially expressed in at least one treatment-time combination in our dataset (Figure 4A).

**Figure 4.**
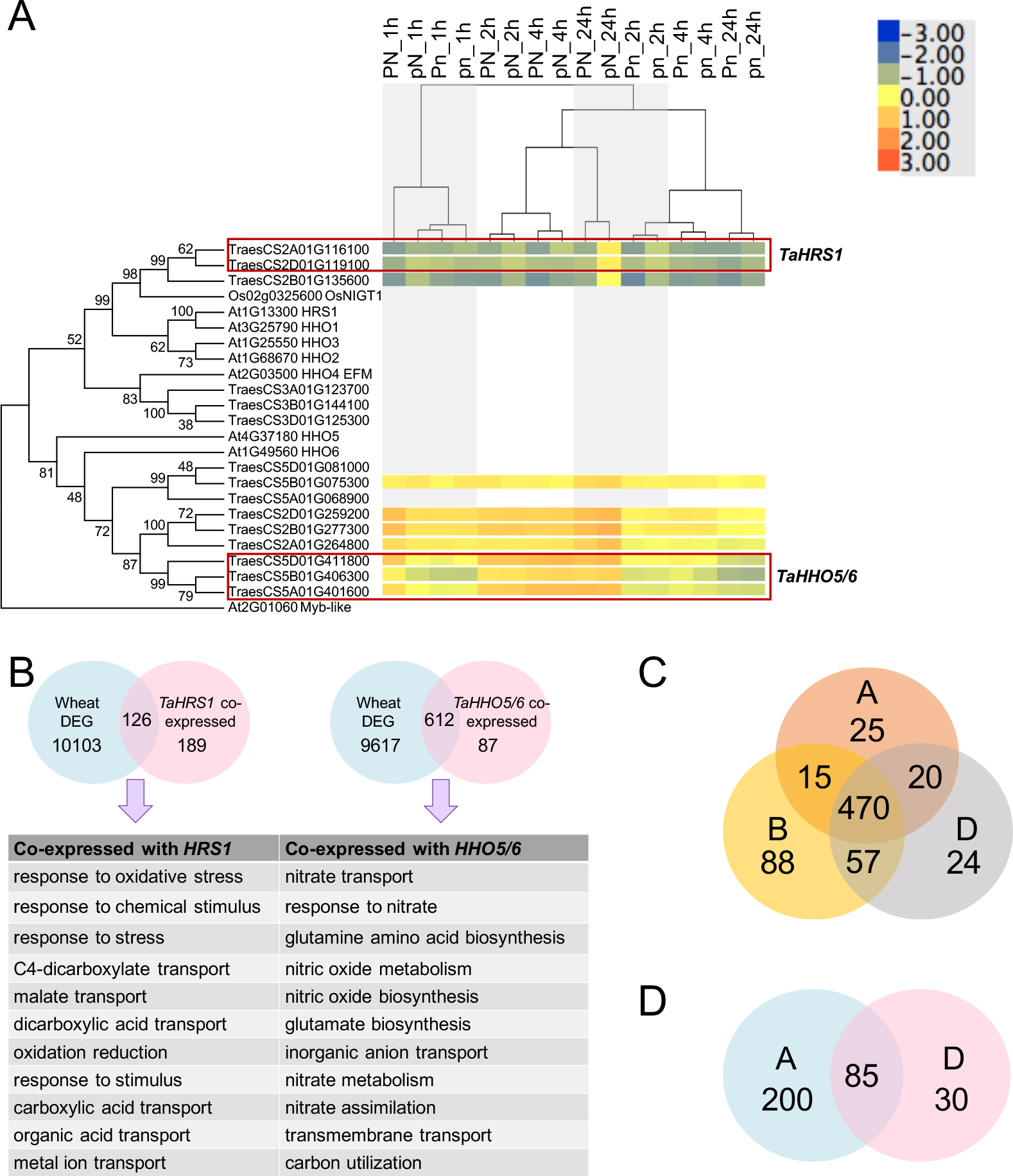
Functional divergence of *HRS1/HHO* gene family members in wheat. **(A)** Maximum likelihood-based (JTT matrix-based model) gene tree for the *HRS1/HHO* gene family members from rice, Arabidopsis and wheat along with gene expression patterns of the wheat *HRS1/HHO* genes; genes in red boxes were assigned to co-expressed modules in the co-expression network. The percentage of trees in which the associated genes clustered together is shown next to the branches. **(B)** Functional differences, determined by GO term enrichment, between genes that are co-expressed with *TaHRS1* or *TaHHO5/6* and are also differentially expressed in response to N/P. **(C)** The overlap of the genes that are co-expressed with wheat *TaHHO5/6* orthologues in a homeologue-specific context - the 699 genes that were co-expressed with *TaHHO5/6* were further grouped based on which homeologue of *TaHHO5/6* they were co-expressed with; 470 (67%) of co-expressed genes were shared by the *TaHHO5/6* triad, **(D)** The overlap of the genes that are co-expressed with wheat *TaHRS1* orthologues in a homeologue-specific context: of the 315 genes that were co-expressed with *TaHRS1* only 27% of co-expressed genes were shared between *TaHRS1_2A* and *TaHRS1_2D*.

Interestingly, the three wheat *HRS1* (*TaHRS1*) genes (*TraesCS2A01G116100*, *TraesCS2B01G135600, TraesCS2D01G119100*) showed up-regulation only at 24 h in response to nitrate supply in the absence of phosphate. This delayed transcriptional response was unexpected of *TaHRS1*, as previous studies on rice (Sawaki *et al.* 2013) and Arabidopsis (Krouk *et al.* 2010) have reported *HRS1/NIGT1* to be up-regulated within 1 h of nitrate supply. However, the seven wheat genes that are closely related to *AtHHO5/HHO6* (*TaHHO5/6* genes) (*TraesCS5B01G075300*, *TraesCS2D01G259200*, *TraesCS2B01G277300*, *TraesCS2A01G264800*, *TraesCS5D01G411800*, *TraesCS5B01G406300*, *TraesCS5A01G401600*) showed up-regulation upon nitrate supply at early time points, and remained induced for the rest of the 24 h of treatment (Figure 4A). Specifically, all *TaHHO5/6* genes but *TraesCS5B01G075300*, were up-regulated by PN treatment at 1 h, and remained up-regulated in PN and pN conditions at later time points. *TraesCS5B01G075300* was up-regulated in PN and pN treatments from 2-24 h (Table S8). This pattern of gene expression indicated that N regulation of the wheat *HRS1/HHO* gene family may have functionally diverged from that of Arabidopsis and rice, and, consequently, the downstream genetic targets of *TaHRS1* and *TaHHO5/6* may also be regulated in a different manner.

It has been estimated that a plant gene can be directly regulated by 6-40 transcription factors (Pal *et al.* 2017). We therefore tested the hypothesis that *TaHRS1* and *TaHHO5/6* differ in their transcriptional responses to N at least in partly due to having different transcription factor binding motifs (TFBM) present within their promoters. We used MEME (Bailey *et al.* 2009) to identify enriched promoter motifs followed by TOMTOM (Khan *et al.* 2018) TFBM comparison analysis of DNA regions 1 kb upstream from transcription start site and identified 11 plant TFBM within the promoter of *TaHRS1* genes (*TraesCS2A01G116100*, *TraesCS2B01G135600, TraesCS2D01G119100*) (Table S7). These 11 TFBM included KANADI, ERF10, homeodomain-like superfamily proteins and Myb-like transcription factors. Except ERF10, the 10 transcription factors predicted to bind to *TaHRS1* promoters all belong to the G2-like transcription factor family (Table S7). On the other hand, the same analysis of the nine *TaHHO5/6* gene promoters (*TraesCS5B01G075300*, *TraesCS2D01G259200*, *TraesCS2B01G277300*, *TraesCS2A01G264800*, *TraesCS5D01G411800*, *TraesCS5B01G406300*, *TraesCS5A01G401600, TraesCS5D01G081000, TraesCS5A01G068900*) identified 118 plant TFBM (Table S7). Among these 118 TFBM, Dof, MADS box and G2-like TFBMs were the most abundant types (Figure S5C, Table S7). All but two G2-like TFBM identified in *TaHRS1* promoters were also present in *TaHHO5/6* promoters (Table S7). The larger diversity of TFBM within the promoters of the *TaHHO5/6* genes may result in increased responsiveness to a wider array of signals. On the other hand, apparent lack of similar level of TFBM diversity within *TaHRS1* promoters may also be due to failure to properly identify wheat-specific TFBM using currently available tools and database of plant TFBM.

In order to identify the regulators of *TaHRS1* and *TaHHO6* we used a different approach where we attempted to infer regulatory networks from gene expression data as implemented in GENIE3 (Huynh-Thu *et al.* 2010). Using this tree-based ensemble approach on the gene expression matrix containing 8300 genes (top 10% genes by expression variance, see above) and the 389 transcription factors (as annotated by the (International Wheat Genome Sequencing Consortium (IWGSC) *et al.* 2018)) present within the 8300 genes, 22 predicted regulators were found to be regulating both members of the homeologue duplet of *TaHRS1* (*TraesCS2A01G116100*, *TraesCS2D01G119100*) (Table S6). Similarly, 29 regulators were commonly predicted for the homeologue triad of *TaHHO5/6* (*TraesCS5D01G411800*, *TraesCS5B01G406300* and *TraesCS5A01G401600*) (Table S6). These genes we refer to as “common regulators” herein. While the majority (10 regulators, Fisher’s exact test *P* = 1.7e-31) of the predicted common regulators were shared between homeologues for *TaHRS1* or *TaHHO6* (Table S6), there were genes that regulated one of the *TaHRS1* or *TaHHO6* homeologues, but not all (Figure 6). Interestingly, the predicted common regulators of *TaHRS1* and *TaHHO6* included wheat homologues of *AtNLP7* and *AtLBD37*. In Arabidopsis, NLP7 is a direct transcriptional regulator of *HRS1/HHO* family members in response to nitrate (Medici *et al.* 2015; Marchive *et al.* 2013) while the role of LBD37 in *HRS1/HHO* regulation remains unclear (Kiba *et al*., 2018; Rubin *et al.* 2009), even though its involvement in regulation of early nitrate responses has been confirmed (Rubin *et al.* 2009; Varala *et al.* 2018), suggesting that LBD37 may act upstream of at least some *HRS1/HHO* family members. Altogether, our results suggest that *TaHRS1/HHO* also require NLP7 and LBD37, while the unique regulators of *TaHRS1* and *TaHHO5/6* possibly contribute to differences in the downstream activity of *TaHRS1* and *TaHHO5/6*.

We further investigated this apparent functional divergence within the wheat *HRS1/HHO* gene family using co-expression network analysis. Out of the 15 *TaHRS1/HHO* homologues, only five genes (*TraesCS2A01G116100*, *TraesCS2D01G119100, TraesCS5D01G411800*, *TraesCS5B01G406300* and *TraesCS5A01G401600*) showed significant co-expression correlation and were assigned to gene co-expression modules (Figure 4A). *TraesCS2A01G116100* and *TraesCS2D01G119100*, which are closely related to *OsNIGT1* (*AtHRS1*), were assigned to turquoise and black modules respectively. The *TaHHO5/6* genes (*TraesCS5D01G411800*, *TraesCS5B01G406300* and *TraesCS5A01G401600)* were assigned to the brown co-expression module. Since *TraesCS2A01G116100* was assigned to the turquoise module, we included the turquoise module in our co-expression sub-network. The GO term enrichment analysis identified nicotianamine metabolism and biogenic amine metabolism processes enriched in the black module (containing the *TaHRS1* gene *TraesCS2D01G119100*), while processes related to oxidative stress were enriched in the turquoise module (containing the *TaHRS1* gene *TraesCS2A01G116100*) (Table S4). As noted above, the brown module, containing the three *TaHHO5/6* homeologues, showed statistically significant enrichment of the GO terms of nitrogen signaling and/or transport (Figure 3B, Table S4).

Furthermore, when we compared the wheat *HRS1* and *HHO5/6* co-expressed genes to the list of differentially expressed genes, 612 out of 699 *HHO5/6* co-expressed genes (88%) were found differentially expressed in response to N/P treatments. The GO term enrichment analysis of these 612 genes identified over-represented processes related to nitrate signaling and metabolism (Figure 4B). On the other hand, the 126 (40%) genes at the intersect of genes co-expressed with *TaHRS1* and the differentially expressed genes do not show any enrichment for processes related to nitrate signaling and metabolism (Figure 4B). These results together with our gene expression profiling results support our hypothesis that the N/P responses in wheat are more likely to be mediated by the *TaHHO5/6* gene regulatory network(s) than the *TaHRS1* one(s).

In order to identify candidate direct TaHRS1- or TaHHO5/6-target gene interactions, we considered the GENIE3 predictions. We then considered the putative common targets of *TraesCS2A01G116100* and *TraesCS2D01G119100* (*TaHRS1*) predicted by the GENIE3 analysis, and found that a significant portion of 205 predicted targets (65%, 133 targets, Fisher’s exact test *P* = 3.6e-85) were also differentially expressed, while only a small proportion of them (14 genes, 7%, Fisher’s exact test *P* = 1.7e-15) were shared with the *TaHRS1* co-expressed genes (Table S6, Figure S5A). Moreover, most of the predicted targets of *TaHRS1* showed similar expression dynamics as *TaHRS1*, whose marked up-regulation was observed only at 24 h in response to pN treatment. Despite this, the GO terms enrichment analysis of the 205 predicted *HRS1* target genes also showed enrichment for GO terms “nicotianamine metabolism” and “cellular amine metabolic process”. This further supports the notion that *TaHRS1* genes may be regulating a class of genes which are functionally different from their counterparts in Arabidopsis. Nicotianamine metabolism plays a crucial role in metal ions transport (Fe and Zn) in plants (Schuler *et al.* 2012; Takahashi *et al.* 2003; Hofmann 2012). Nicotianamine has been considered a metal ion chelator that enables proper distribution of metal ions among source and sink tissues (Schuler *et al.* 2012; Conte and Walker 2011), while contributing to prevent metal ion toxicity (Wiren N *et al.* 1999). Up-regulation of nicotianamine synthase genes have been reported in response to nitrate supply which has been thought to be a response to facilitate transport of Fe required for synthesis of nitrate assimilatory enzymes such as nitrate reductase and nitrite reductase (R. Wang *et al.* 2003). While nicotianamine synthase gene *NAS3* was induced under P starvation (Bournier *et al.* 2013), *NAS2* was up-regulated under excess P condition (Shukla *et al.* 2017). Therefore, nitrate or phosphate treatments trigger changes in the nicotianamine levels which may lead to precise regulation of metal ion transport. Considering the gene expression dynamics of *TaHRS1* homeologue triad, the GENIE3-predicted targets of *TraesCS2A01G116100*, *TraesCS2D01G119100*, the biological processes enriched in the black and turquoise co-expression modules, and the biological processes enriched in the predicted targets of *TaHRS1*, we propose that *TaHRS1* may be regulating nitrate-dependent metal ion transport in the absence of phosphate.

Similar to the results of the WGCNA analysis, GENIE3 identified a larger set of predicted common gene targets of *TraesCS5D01G411800*, *TraesCS5B01G406300* and *TraesCS5A01G401600* (*TaHHO5/6*) than *TaHRS1* (Table S6) with 754 predicted *TaHHO5/6* targets. While 644 out of 754 predicted targets (85%, Fisher’s exact test *P* = 0) were differentially expressed in at least one treatment-time combination, a much larger proportion (537 genes) of the *TaHHO5/6* predicted targets were also co-expressed with *TaHHO5/6* genes (71%, Fisher’s exact test *P* = 0) (Figure S5B, Table S6). GO term enrichment analysis on these 754 predicted TaHHO5/6 targets revealed that processes such as “response to nitrate”, “nitrate transport”, “glutamate biosynthetic process” were enriched.

This result corroborates our hypothesis of the possible functional convergence in the background of genetic divergence of some of the *TaHRS1/HHO* family members in mediating nitrate transcriptional responses and the regulatory roles these two transcription factors play in wheat P/N responses. Altogether our results suggest that, *TaHRS1* and *AtHRS1* are functionally divergent, while *TaHHO5* and *AtHRS1* show functional convergence while being genetically divergent. These differences may be due to *TaHHO5/6* rapid induction by nitrate that initiates a transcriptional response cascade as a primary response to nitrate supply. Meanwhile, *TaHRS1* is induced later in the time course and regulates secondary processes, such as metal ion transport. However, it should also be noted that these putative transcription factor – target relationships are predicted based on gene expression dynamics and the post-transcriptional level regulation by *TaHRS1/HHO* members may be different.

In line with their divergent regulation and predicted function as transcription factors, the set of genes that are co-expressed with either the two *TaHRS1* or the three *TaHHO5/6* homeoalleles are different, despite their homeologue relationships. For example, *TraesCS2A01G116100* and *TraesCS2D01G119100*, the *TaHRS1* homeoalleles, share 85 co-expressed genes (85/315 or only 27%) (Figure 4D). Meanwhile, the *TaHHO5/6* homeoallele triad (*TraesCS5D01G411800*, *TraesCS5B01G406300* and *TraesCS5A01G401600*) shares 470 co-expressed genes (66% of genes co-expressed with the entire *TaHHO5/6* triad) while another 92 genes (13%) are co-expressed with either two of the three *TaHHO5/6* homeoalleles (Figure 4C). This suggests that presence of active homeoalleles may increase the complexity of wheat gene regulatory networks.

Altogether, these results suggest that *TaHHO5/6* orthologues are functionally more related to *AtHRS1* and *OsNIGT1*(Figure 4B), while *TaHRS1* may have a different role in mediating N/P responses in wheat, in line with its delayed transcriptional responses (Figure 4A).

The *TGA* gene family members are transcription factors characterized by the presence of bZIP_1 signature DNA binding domain and their ability to bind to the TGAGC promoter motif (Lebel *et al.* 1998; Gatz 2013). In Arabidopsis, 10 *TGA* gene family members have been identified (Jakoby *et al.* 2002). *TGA* family members were first identified to be involved in pathogenesis related processes (Subramaniam *et al.* 2001; Fan and Dong 2002), however, *TGA1* and *TGA4* have been shown to regulate root development in response to nitrate (Álvarez *et al.* 2014). None of the other Arabidopsis *TGA* family members have been found to be involved in nitrate signaling/metabolism (Álvarez *et al.* 2014). Our analysis showed that the *TGA* gene family in wheat consists of at least 38 members (as identified through reciprocal BLAST), all of which possess the conserved bZIP_1 DNA binding domain (Figure S3B-C). Out of 9 wheat *TGA* genes (*TraesCS3A01G372400*, *TraesCS3D01G365200*, *TraesCS3B01G404800*, *TraesCS1A01G276600*, *TraesCS1B01G285800*, *TraesCS1D01G276100*, *TraesCS4A01G183400*, *TraesCS4B01G135000* and *TraesCS4D01G129900*) that form a separate clade with three members of the rice *TGA* family, 8 genes were differentially expressed in at least one treatment-time combination (Figure 5A). Since only *TraesCS1A01G276600*, *TraesCS1B01G285800* and *TraesCS1D01G276100* were assigned to modules in WGCNA analysis and differentially expressed, we focused more on these three genes. Based on sequence homology, *TraesCS1A01G276600*, *TraesCS1B01G285800* and *TraesCS1D01G276100*, (constituting a homeologue triad) were found to be most closely related to *Os05g0443900* (*Liguleless2*), through a reciprocal BLAST (Figure 5A, Figure S4). Since *AtTGA9/10* are the closest Arabidopsis *TGA* family members to *TraesCS1A01G276600*, *TraesCS1B01G285800* and *TraesCS1D01G276100* on the gene tree (Figure 5A, Figure S4), we refer to *TraesCS1A01G276600*, *TraesCS1B01G285800* and *TraesCS1D01G276100* triad as *TaTGA9/10* herein. In contrast to Arabidopsis, we did not detect differential expression of wheat *TGA1/4* homologues (*TraesCS2A01G526300*, *TraesCS2B01G556600*, *TraesCS2D01G529000)*, and they were not assigned to modules in the WGCNA network. However, we detected *TaTGA9/10* orthologues as differentially expressed (up-regulated under PN condition throughout the time course and under pN condition 2-24 h (Table S8)) and co-expressed with nitrate signaling/metabolism related genes in the brown module (Figure 4C). This may suggest possible sub-functionalization of wheat *TGA* members in regulating nitrate signaling/metabolism. Similar to the *TaHHO5/6* homeologue triad, when the *TaTGA9/10* co-expressed genes were considered in a homeologue-specific context, 454 genes out of 517 genes (88% of co-expressed genes with *TaTGA9/10* triad) were co-expressed with all three *TaTGA9/10* genes (Figure 5B). Since the *TaTGA9/10* triad was loosely annotated as “transcription factor like proteins”, our gene co-expression analysis identified putative functions of this homeologue triad in the context of nitrate signaling/metabolism. And our results suggest that in wheat, *TaTGA9/10* triad is functionally convergent with *AtTGA1/4* despite being genetically divergent.

**Figure 5.**
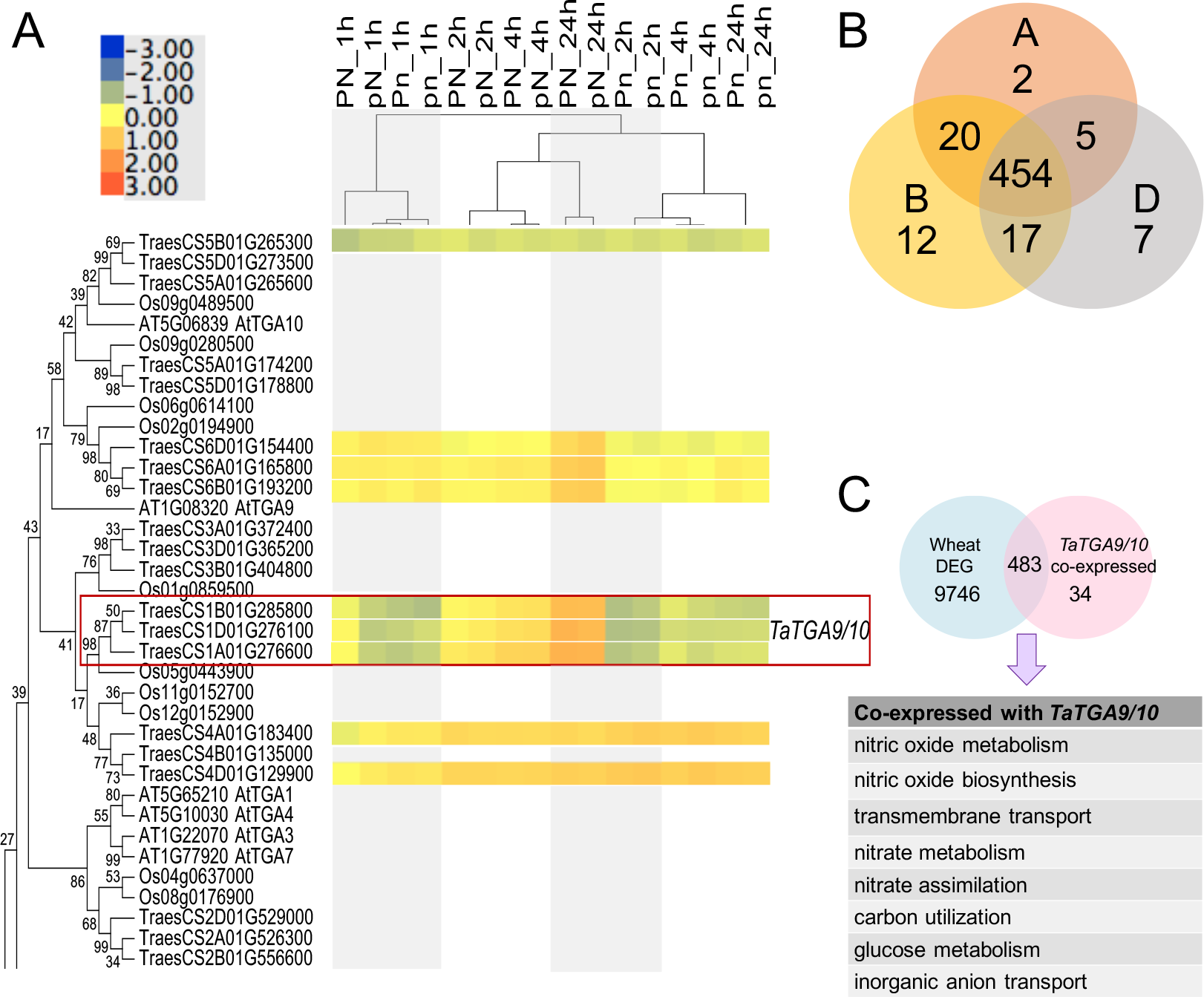
Functional divergence of *TGA* gene family members in wheat. **(A)** Segment of the maximum likelihood based (JTT matrix-based model) gene tree for the *TGA* family members from rice, Arabidopsis and wheat along with gene expression patterns of wheat *TGA* orthologues; genes in red boxes were assigned to co-expressed modules in the coexpression network. The percentage of trees in which the associated genes clustered together is shown next to the branches; full gene tree is in Figure S4. **(B)** The overlap of the genes that are co-expressed with *TaTGA9/10* orthologues in a homeologue-specific context; the 517 genes that were co-expressed with *TaTGA9/10* were further grouped based on which homeologue of *TaTGA9/10* they were co-expressed with; 88% of co-expressed genes were shared by *TaTGA9/10* triad. **(C)** Genes that are both co-expressed with *TaTGA9/10* and differentially expressed are involved in nitrate signaling and metabolism.

Moreover, our GENIE3 analysis predicted 1107 common putative target genes downstream of *TaTGA9/10* genes (Table S6). These are enriched for processes such as “nitrate assimilation”, “malate transport” and “glutamate biosynthetic processes”, with 850 of the 1107 (76.7%, Fisher’s exact *P* = 0) targets also differentially expressed in at least one of the treatment-time combinations, while 447 targets (40.3%, Fisher’s exact *P* = 0) were co-expressed with *TaTGA9/10*. Intriguingly, phosphate transporters (12 genes homologous to *AtPT1* and *AtPHO1*) and SPX domain containing proteins (8 genes homologous to *AtSPX2/3*, *AtPHT5;1*) were also among the predicted TaTGA9/10 targets (Table S6), out of which 4 phosphate transporter genes and 3 SPX domain containing genes were co-expressed as well. Indeed it has been reported that most phosphate transporters and phosphate starvation induced genes, including *SPX1/3/4* and *PHO1*, possess TGAGC motif (Baek *et al.* 2017). Our results suggest that *TaTGA9/10* participates in the nitrate dependent regulation of phosphate starvation responses and transport, and as such, likely plays a role in integrating nitrate and phosphate signals.

Considering the evidence for possible functional divergence of *TaHRS1/HHO* and *TaTGA* gene family members, with *TaHHO5/6* and *TaTGA9/10* being prominent in wheat nitrate signaling/metabolism by regulating hundreds of target genes, we expected that a transcriptional network of *TaHRS1/HHO* and *TaTG9/10* members and their regulators would be important in identifying regulatory mechanisms underpinning *TaHRS1/HHO* and *TaTGA9/10* transcriptional responses to N and P provision. To this end, we used the GENIE predictions to identify putative regulatory relationships between transcriptional regulators of *TaHRS1* homeologue duplet, *TaHHO5/6* homeologue triad, *TaTGA9/10* homeologue triad (Figure 6, Table S9). Indeed, we found that wheat *NLP7* genes (*TraesCS6A01G102400*, *TraesCS6B01G130800* and *TraesCS6D01G091000*) were among regulators shared between *TaHRS1*, *TaHHO5/6* and *TaTGA9/10* highlighting the central regulatory role played by *NLP7* genes in wheat nitrate responses. In addition to wheat *NLP7* genes, *TraesCS6A01G287700* (Dof zinc finger protein homologous to *AT2G37590*) and a homeologue triad (*TraesCS2A01G488200*, *TraesCS2B01G515800* and *TraesCS2D01G488500*) of a MYB-like transcription family protein homologous to *AT5G06800*, a MYB-HTH transcriptional regulator family protein, were also shared regulators of *TaHRS1*, *TaHHO5/6* and *TaTGA9/10* (Figure 6, Table S9), suggesting that these transcription factors play a role in regulating wheat nitrate responses. While *AT2G37590* and *AT5G06800* have not been characterized in the context of nitrate regulation, gene co-expression data from Arabidopsis suggest that *AT5G06800* is co-expressed with *AtHRS1* (Obayashi *et al.* 2018). Moreover, the targets of wheat *LBD37* genes (*TraesCS4B01G078800*, *TraesCS2B01G212400*, *TraesCS4D01G077600* and *TraesCS4A01G236200*) also included *TaHRS1*, *TaHHO5/6* and *TaTGA9/10* (Figure 6, Table S9). These results are consistent with observations made about NLP7 (Medici *et al.* 2015; Marchive *et al.* 2013) and LBD37 (Rubin *et al.* 2009; Varala *et al.* 2018) regulatory roles in Arabidopsis. *TaHHO5/6* and *TaTGA9/10* expression dynamics were also consistent with their being regulated by each other, as predicted by the GENIE3 analysis. However, the exact nature of this cross-regulation remains unresolved due to a lack of fine scale temporal resolution in our dataset.

**Figure 6.**
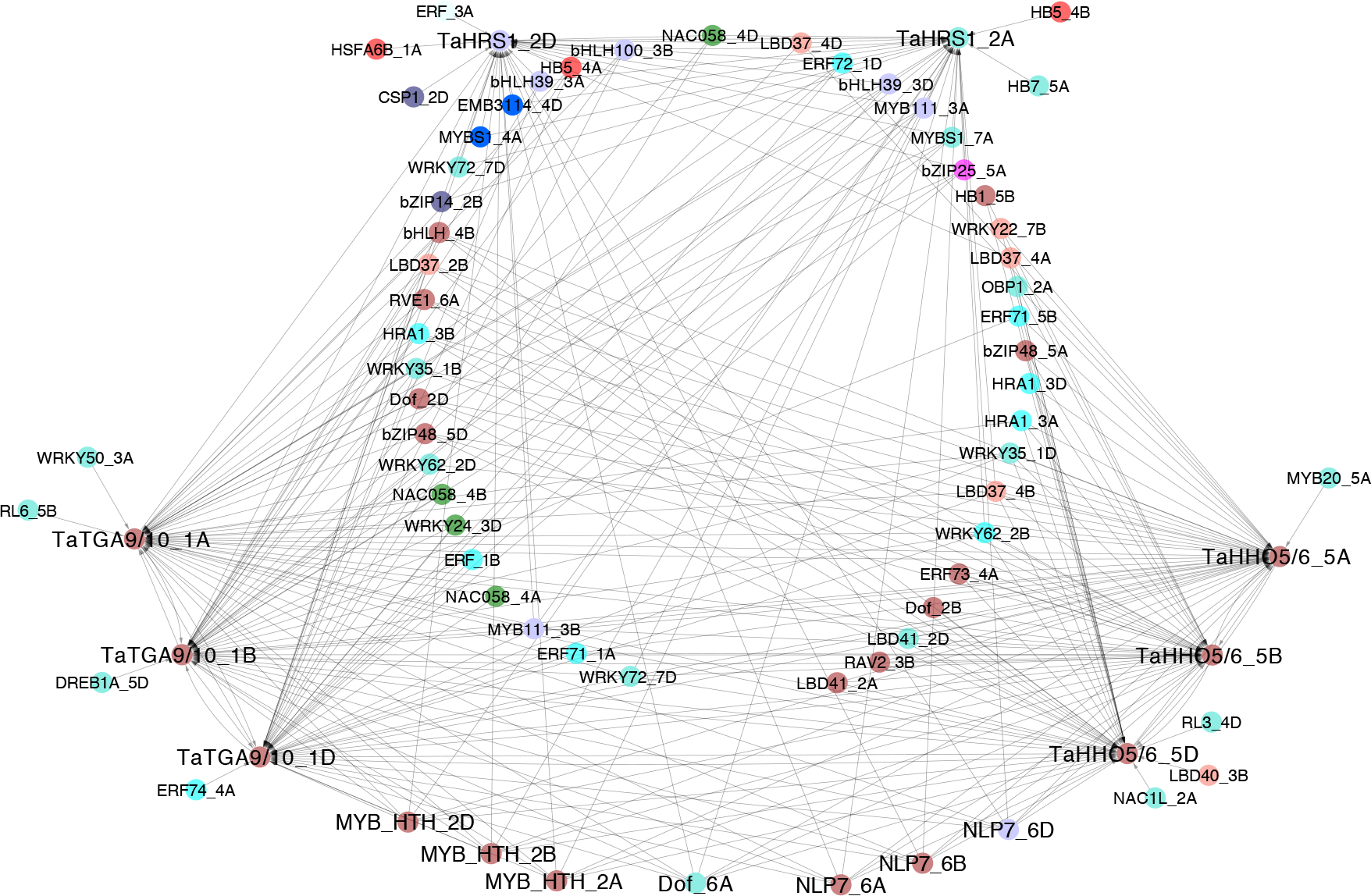
Predicted relationships between *TaHRS1*, *TaHHO5/6*, *TaTGA9/10* and their regulators. The node colors correspond to the module color on WGCNA network, except genes belonging to black module are represented in light purple for clarity. Edges represent a weight measure >0.005 as calculated by GENIE3. The nodes are named based on their sequence homology to Arabidopsis.

In our transcriptional regulatory network, we could also identify regulators that were predicted to be regulating only *TaHRS1* or *TaHHO5/6* or *TaTGA9/10* (Figure 6, Table S9). For example, wheat genes homologous to *AtLBD41* and *AtRAV2* were predicted to be specifically regulating only the *TaHHO5/6* triad, while genes homologous to *AtNAC058* and *AtWRKY24* were predicted to be specifically regulating *TaTGA9/10* triad (Figure 6, Table S9). Although these regulatory relationships are predicted based on gene expression and therefore any post-transcriptional/translational effects are likely missed, our results suggest that the regulatory roles played by genes such as *NLP7* and *LBD37* are conserved in wheat, while identifying candidate genes, such as *TraesCS2A01G488200*, *TraesCS2B01G515800* and *TraesCS2D01G488500*, which may play a central role in regulating wheat nitrate Plants sense multiple nutritional signals and show coordinated responses to these signals as suggested by the emerging evidence for crosstalk between nutrient signaling mechanisms such as those between nitrate and phosphate signaling (Kellermeier *et al.* 2014; Medici *et al.* 2015). Hence, studying the combined effects exerted by nitrate and phosphate on the transcriptional response within wheat roots identified the synergistic effects of N and P supply on gene expression as early as 1 h post-treatment, while also identifying other, mutually independent transcriptional responses in wheat. Using recent wheat genome annotation, we identified instances of likely functional divergence in transcription factor gene families prominent in nitrate signaling and metabolism. Analysis of gene co-expression networks identified gene modules that are involved in wheat nitrate signaling and metabolism which begins to unravel the complex and intricate nutrient signaling components in wheat and will be helpful in prioritizing candidate genes to increase nutrient use efficiency in wheat.

## Materials and methods

### Plant growth and treatment

Surface sterilized wheat seeds (variety Chinese spring) were pre-imbibed at 4 °C overnight and transferred on to moistened rock wool plugs in a hydroponics system and the plants were grown at 24 °C in a growth chamber at 16 h photoperiod. The seedlings were grown in water for 11 days before treating them with nutrients. After 11 days, water was replaced with nutrient solutions of varying nitrate and phosphate concentrations. Composition of the nutrient solutions is available in Table S10. Briefly, the plants were subjected to 0.5 mM (P-sufficient) or 0 mM (P-deficient) phosphate in the form of KH2PO4 and 2 mM (N-sufficient) or 0 mM (N-deficient) nitrate in the form of Ca(NO3)2. The nutrient solution was changed once in every three days and for RNA extraction, total roots were harvested after 1 h, 2 h, 4 h and 24 h of transferring to the nutrient solution, snap-frozen in liquid nitrogen and stored at −80 °C. The plant root measurements were taken after 7 days of treatment. Four – six of 18-da old wheat seedlings that were grown in nutrient solutions were used for *in planta* nutrient content measurements. Nitrogen content was analyzed by combustion. Phosphorus content was analyzed by ICPOES on samples open-vessel digested using 5:1 mixture of nitric and perchloric acids.

### RNA extraction and library preparation

Samples were collected in triplicate per treatment-time point combination and each sample contained root tissue from four - six plants. Total RNA was extracted from 48 samples using TRI-reagent (SIGMA-ALDRICH) according to manufacturer’s instruction. The RNA samples were treated with DNase I (NEW ENGLAND BIOLABS) and poly-A RNA selection was carried out using Dynabeads mRNA purification kit (THERMOFISHER SCIENTIFIC) according to manufacturer’s instructions. RNA-Seq library preparation was carried out using NEBNext Ultra RNAlibrary prep kit for Illumina (NEW ENGLAND BIOLABS) and 48 RNA-Seq libraries were sequenced using 75×2 setting on Illumina HiSeq2500 platform to obtain at least 20 million 75 bp paired end reads per sample (Table S1).

### RNA-Seq data analysis

#### Data pre-processing

Each RNA-Seq library had more than 20 million reads and initial quality control was carried out using FastQC package (Andrews, 2010). The libraries were pre-processed to remove adapter sequences and low quality reads using Trimmomatic (Bolger *et al.* 2014). Wheat transcript file was generated through RSEM (B. Li and Dewey 2011) using IWGSC wheat RefSeqv1.0 genome assembly (161010_Chinese_Spring_v1.0_pseudomolecules.fasta) and annotation (iwgsc_refseqv1.0_HighConf_2017Mar13.gff3) (International Wheat Genome Sequencing Consortium (IWGSC) *et al.* 2018). Next, the transcripts were used to take the read counts for each transcript/gene using Salmon – a quasi-mapping based algorithm (Patro *et al.* 2017). The samples had average mapping percentage of 68% (Table S1).

#### Differential gene expression analysis

Differential gene expression analysis was performed using DESeq2 version 1.18.1 (Love *et al.* 2014) using a design formula (~time + treatment + treatment:time) that takes into account the contrast between two treatment levels throughout the time course as well at a FDR<0.05. A complete list of differentially expressed genes at each treatment-time point along with the log fold change and adjusted P values is available in Table S8. The differentially expressed gene lists were compared using BioVenn (Hulsen *et al.* 2008). For the non-redundant list of differentially expressed genes, the z-score normalized variance stabilized expression values were used to generate hierarchical clustering based heat map using pheatmap R package (Kolde, 2013).

#### GO term enrichment analysis

GO term analysis was carried out using BinGO app for Cytoscape (Maere *et al.* 2005) for a custom made wheat GO term annotation database for IWGSCRefseqv1.0 (International Wheat Genome Sequencing Consortium (IWGSC) *et al.* 2018). The most terminal GO terms that are enriched in the hierarchy were considered for further analyses. Fold enrichment was calculated as (x/X)/(n/N) where x is the number of genes (from the list) assigned to a GO category, X is the total number of genes (in the gene list) that were assigned to any GO category, n is the number of genes (in the genome) assigned to the considered GO category and N is the total number of genes (in the genome) that were assigned to GO categories. Over-represented GO terms at FDR < 0.05 were considered for further analyses.

#### Gene co-expression network analysis

Variance stabilized count data of the expressed genes were considered and the genes were ranked by variance of gene expression across all conditions. The top 8300 (top 10%) genes were selected for gene co-expression network analysis using WGCNA package version 1.64.1 (Langfelder and Horvath 2008). Signed network with soft-thresholding power of 12 was generated. The co-expression network was visualized in Cytoscape version 3.5.1 (Shannon *et al.* 2003). GO term enrichment analysis for the selected modules was carried out using BinGO app for Cytoscape (Maere *et al.* 2005).

### Gene tree construction

The wheat orthologues for *HRS1/HHO* and *TGA* gene family members were identified by a reciprocal tBLASTx search with a E value cutoff of 10^−5^. The amino acid sequences of the *HRS1/HHO* and *TGA* gene family orthologues from Arabidopsis, rice and wheat were aligned using ClustalW (Thompson *et al.* 1994) and phylogenetic tree construction was done using maximum likelihood method (JTT matrix-based model) (Jones *et al.* 1992) for 1000 bootstrap replicates using the MEGA6.06 (Tamura *et al.* 2013).

### Promoter motif analysis

For the considered genes in wheat, 1000 bp upstream region nucleotide sequences were extracted and used for MEME algorithm for motif discovery (version 5.0.1) (Bailey *et al.* 2009) with default parameters and −nmotifs 30. The results from MEME motif search were then used to run TOMTOM algorithm for motif comparison (version 5.0.1) (Bailey *et al.* 2009) against JASPAR non-redundant core plant PFM database (release 7, 2018) (Khan *et al.* 2018).

To search for the conserved domains in putative wheat TGA family members, the amino acid sequences were extracted for the TGA family members and these sequences were used to perform MEME algorithm for motif discovery (version 5.0.1) (Bailey *et al.* 2009) with protein sequence alphabet.

### GENIE3 analysis

Variance stabilized count data of the expressed genes were considered and the genes were ranked by variance of gene expression across all conditions. The top 8300 (top 10%) genes were selected for GENIE3 analysis using GENIE3 version 1.4.0 (Huynh-Thu *et al.* 2010). There were 389 genes that were annotated as transcription factors or transcription factor like proteins with evidence from GO-, Pfam- or Interpro-IDs among the selected 8300 genes. Random forest method was used to infer the network with default paramaters with gene expression matrix for 8300 genes and a list of transcription factor genes as the input. The regulatory relationships that had a weight measure >0.005 were considered for downstream analyses which included GO term enrichment analysis (as described above) for predicted target genes of *TaHRS1*, *TaHHO5/6* and *TaTGA9/10* and gene regulatory network visualization using Cytoscape version 3.5.1 (Shannon *et al.* 2003).

### Gene list overlaps

Fisher’s exact test for overlap between gene lists was performed through GeneOverlap R package (Shen and Senai, 2013). Overlap was considered significant if the adjusted *P* value < 0.05.

## Supporting information

Supplemental data for Dissanayake et al

## Author Contributions

M.T. and I.D. conceived the project. I.D. carried out the experiments, collected the samples and prepared the RNA-Seq libraries. I.D. carried out data analyses with contributions from J.R-M. I.D., J.R-M., S.B. and M.T. interpreted the data. I.D. and M.T. wrote the manuscript with input from S.B. and J.R-M.

## Supplementary Figures

**Figure S1.**
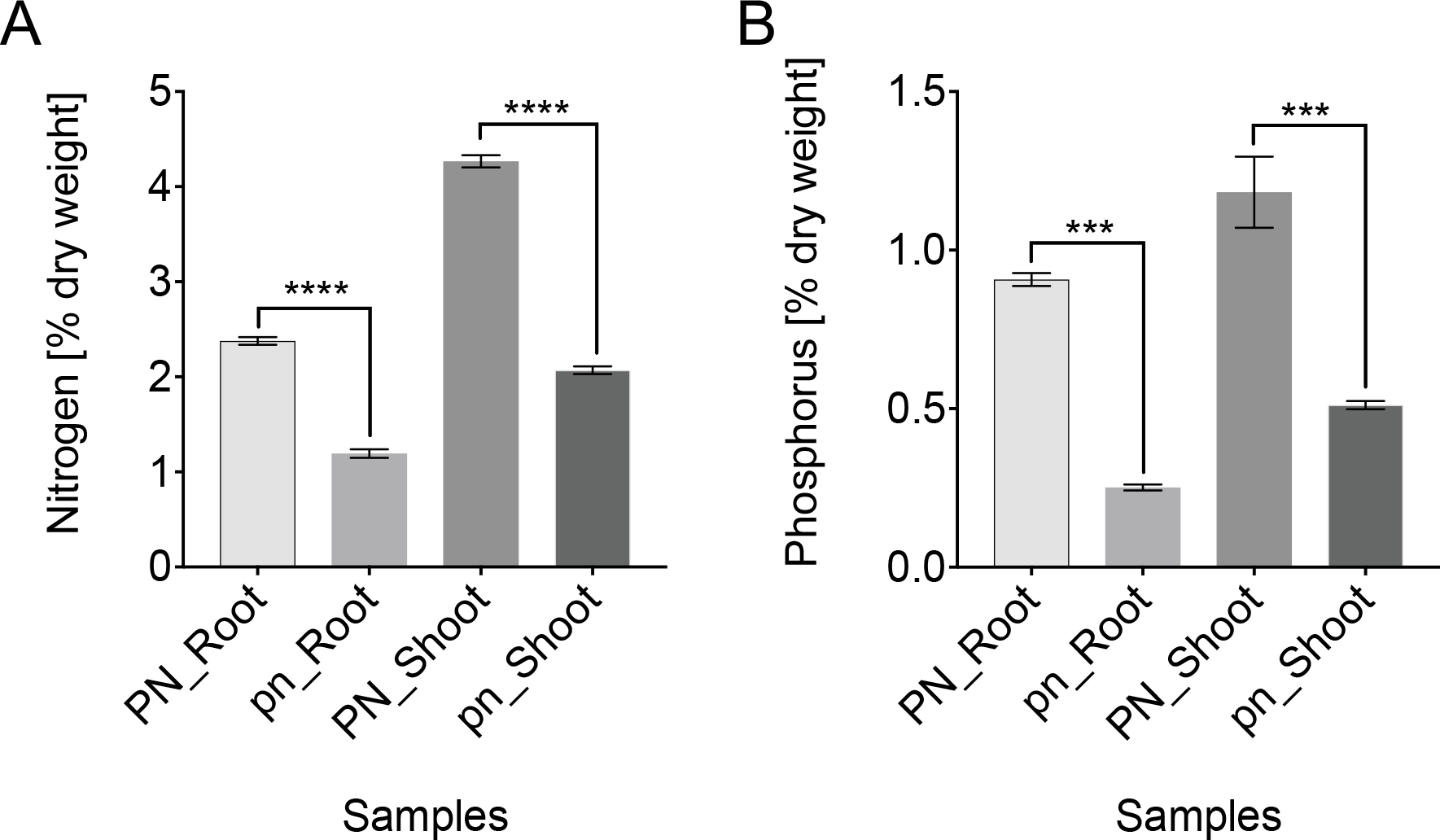
**(A)** Nitrogen and **(B)** Phosphorus content (as a percentage of dry weight) in 11 days old wheat roots and shoots that were grown in deionized water (pn) or full strength nutrient solution (PN: 2 mM nitrate and 0.5 mM phosphate) (n=3, error bars indicate standard error of mean, significance *P*<0.05).

**Figure S2.**
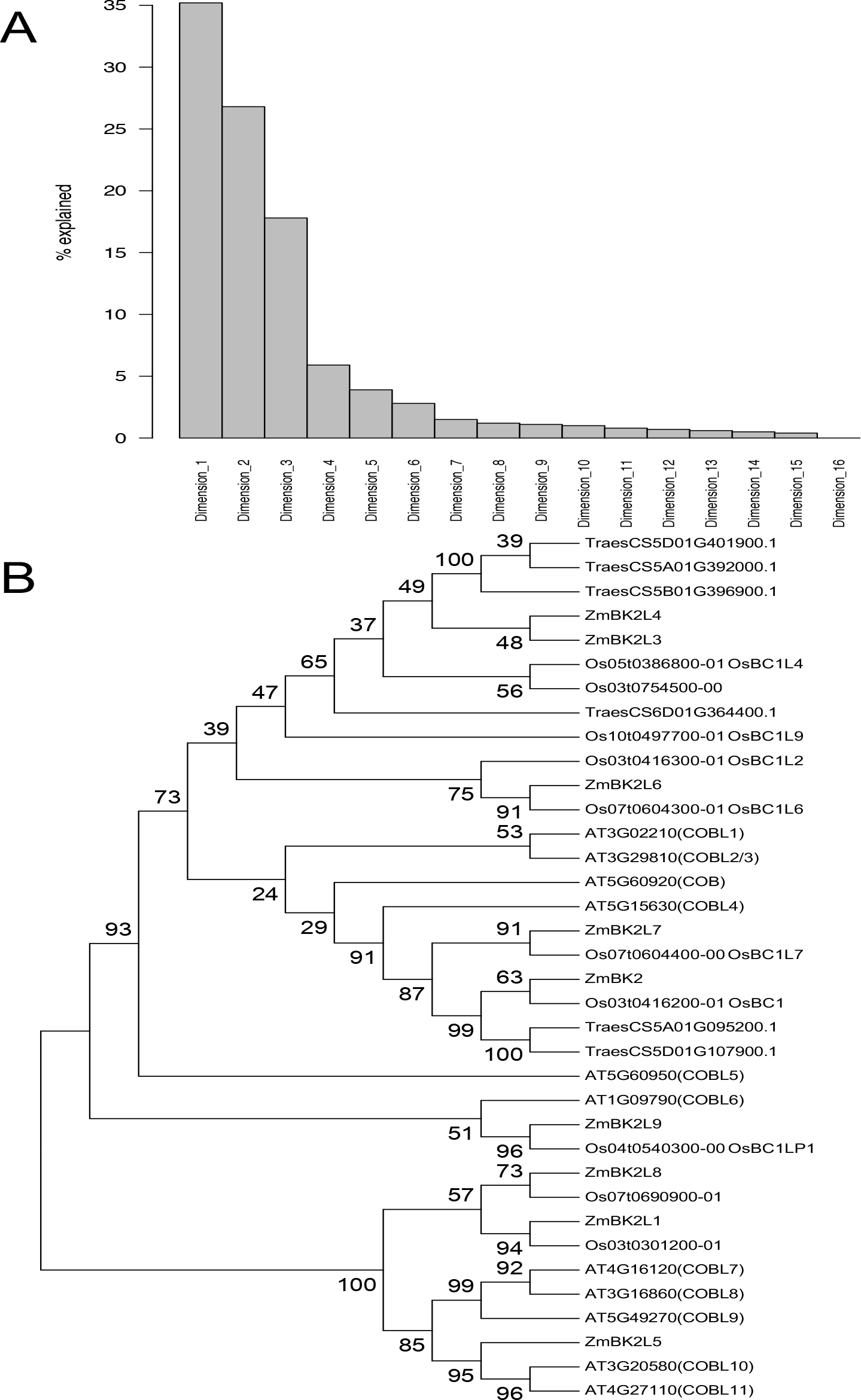
**(A)** Percentage of variance of the dataset explained by each dimension in multidimensional scaling plot (MDS plot) where number of dimensions was arbitrarily set to 16. **(B)** Maximum likelihood based (JTT matrix-based model) phylogenetic tree for the *COBRA* family members from rice, maize, Arabidopsis and wheat. The percentage of trees in which the associated taxa clustered together is shown next to the branches.

**Figure S3.**
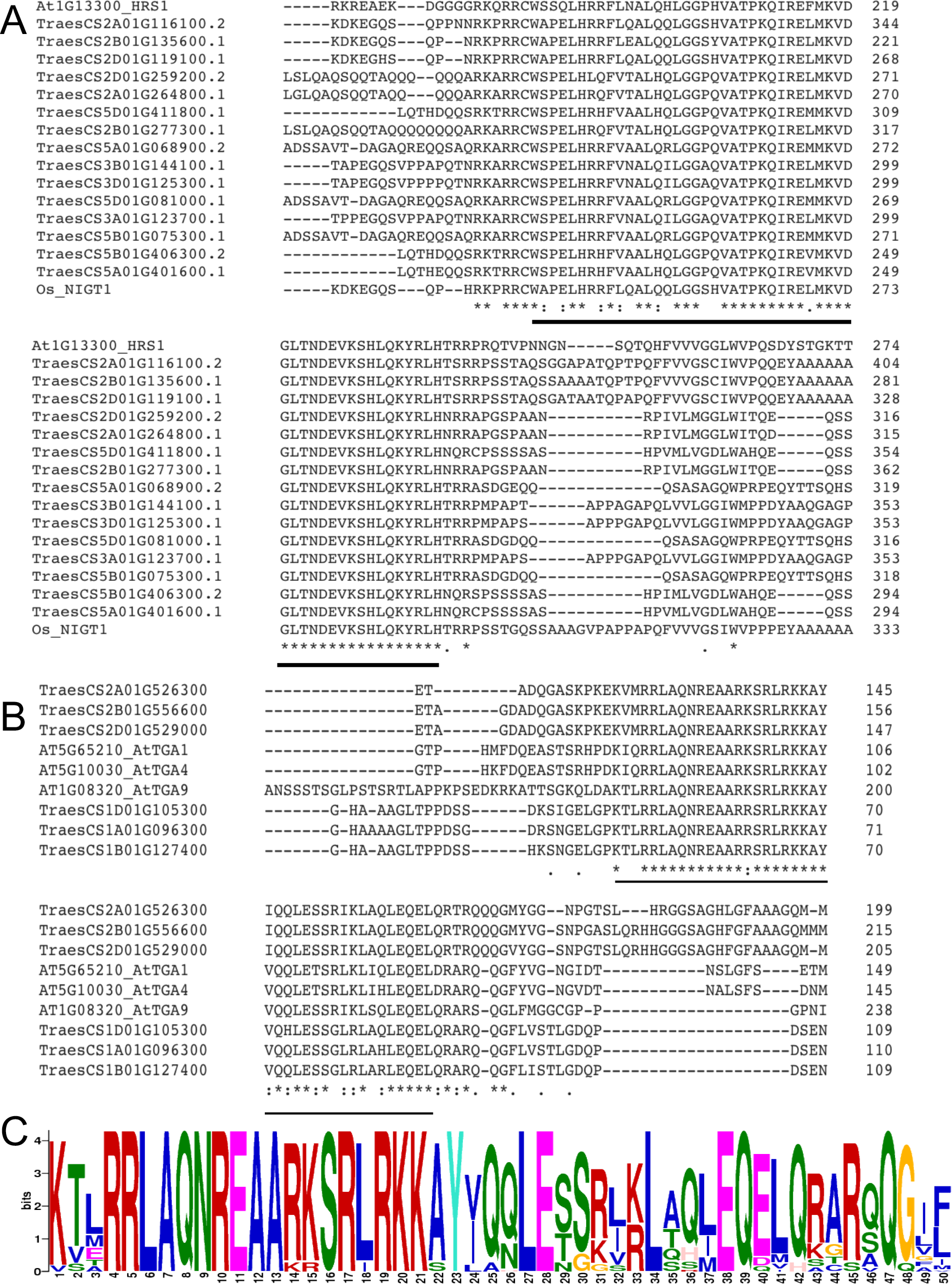
**(A)** The amino acid sequence alignment for AtHRS1, OsNIGT1 and members of wheat HRS1/HHO family; the conserved G2-like DNA binding domain is indicated with the black bar. **(B)** The amino acid sequence alignment for AtTGA1, AtTGA4, AtTGA9 and wheat orthologues for TGA1/4 and TGA9/10; the conserved bZIP_1 signature domain is indicated with the black bar. **(C)** The most enriched motif in putative *TaTGA* family members.

**Figure S4.**
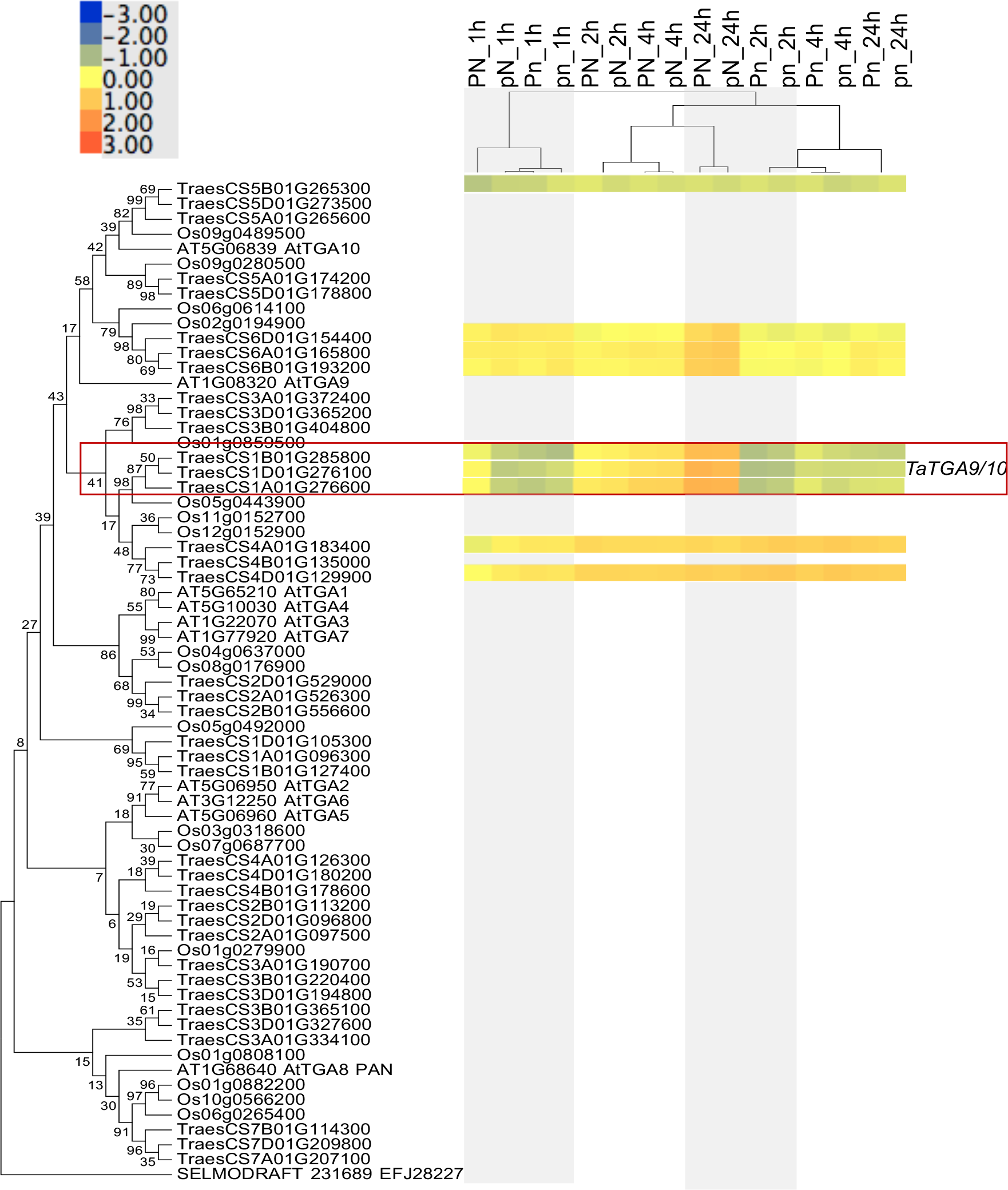
Maximum likelihood based (JTT matrix-based model) phylogenetic tree for the *TGA* family orthologues from rice, Arabidopsis and wheat along with gene expression patterns of wheat orthologues; genes in red boxes were assigned to brown module in WGCNA.

**Figure S5.**
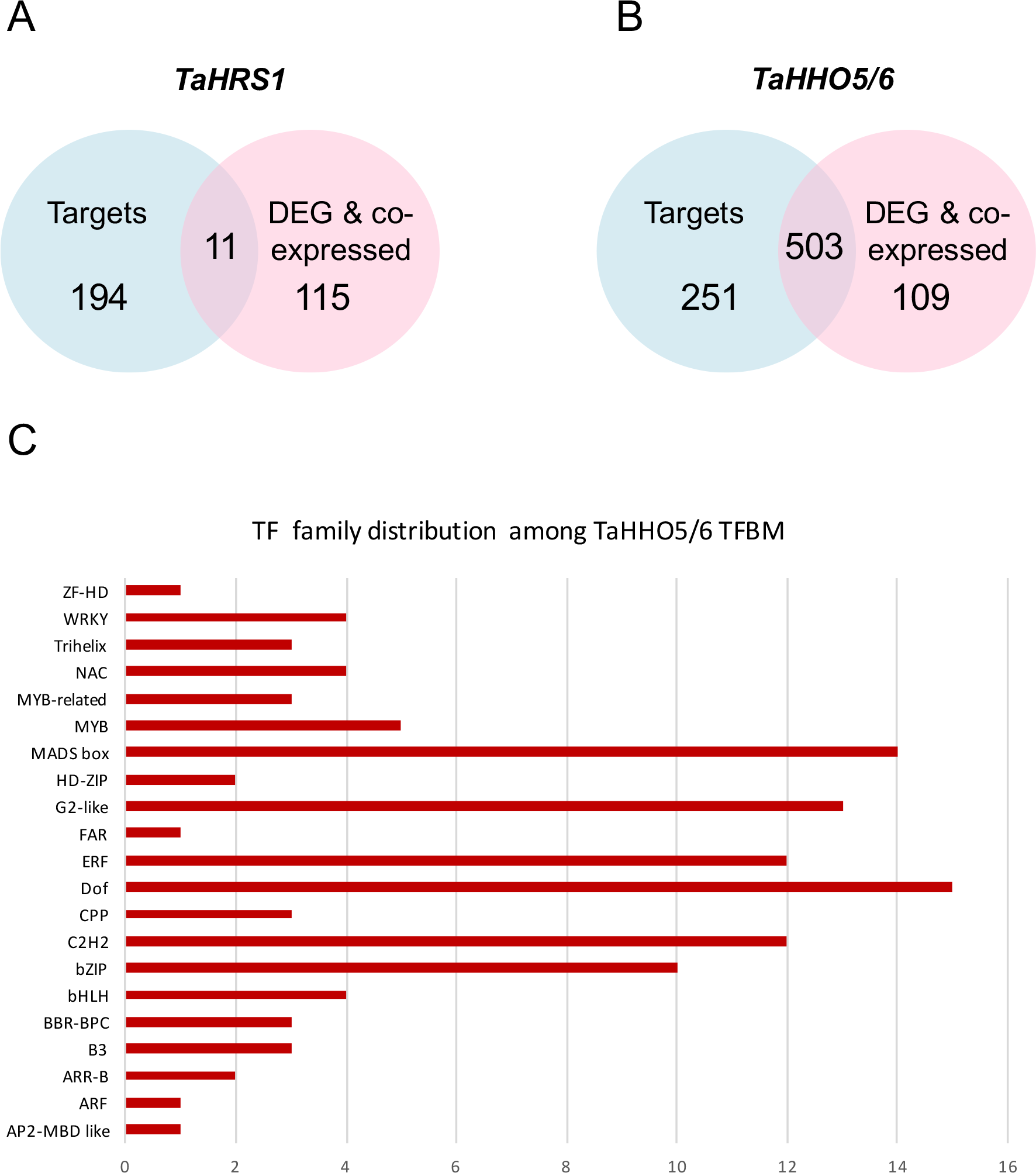
The overlap between targets predicted by GENIE3 with genes that are both differentially expressed and co-expressed with, **(A)** *TaHRS1* or **(B**) *TaHHO5/6*. **(C)** Distribution of TaHHO5/6 transcription factor binding motifs (TFBM) based on transcription factor (TF) family.

## References

Abel, S. and Theologis, A. (1996). Early genes and auxin action. PLANT PHYSIOLOGY 111:9–17.

Álvarez, J.M., Riveras, E., Vidal, E.A., Gras, D.E., Contreras-López, O., Tamayo, K.P., Aceituno, F., et al. (2014). Systems approach identifies TGA1 and TGA4 transcription factors as important regulatory components of the nitrate response of Arabidopsis thalianaroots. The Plant Journal 80:1–13.

Baek, D., Chun, H.J., Yun, D.-J. and Kim, M.C. (2017). Cross-talk between Phosphate Starvation and Other Environmental Stress Signaling Pathways in Plants. Molecules and cells 40:697–705.

Bailey, T.L., Boden, M., Buske, F.A., Frith, M., Grant, C.E., Clementi, L., Ren, J., et al. (2009). MEME SUITE: tools for motif discovery and searching. Nucleic acids research 37:W202–8.

Balzergue, C., Dartevelle, T., Godon, C., Laugier, E., Meisrimler, C., Teulon, J.-M., Creff, A., et al. (2017). Low phosphate activates STOP1-ALMT1 to rapidly inhibit root cell elongation. Nature Communications 8:15300.

Banf, M. and Rhee, S.Y. (2017). Computational inference of gene regulatory networks: Approaches, limitations and opportunities. BBA - Gene Regulatory Mechanisms 1860:41–52.

Beadle, N.C.W. (1953). The Edaphic Factor in Plant Ecology With a Special Note on Soil Phosphates. Ecology 34:426–428.

Benfey, P.N. and Scheres, B. (2000). Root development. Current Biology 10:R813–5.

Bieleski, R.L. (1973). Phosphate Pools, Phosphate Transport, and Phosphate Availability. Annual Review of Plant Physiology 24:225–252.

Bolger, A.M., Lohse, M. and Usadel, B. (2014). Trimmomatic: a flexible trimmer for Illumina sequence data. Bioinformatics (Oxford, England) 30:2114–2120.

Bournier, M., Tissot, N., Mari, S., Boucherez, J., Lacombe, E., Briat, J.-F. and Gaymard, F. (2013). Arabidopsis ferritin 1 (AtFer1) gene regulation by the phosphate starvation response 1 (AtPHR1) transcription factor reveals a direct molecular link between iron and phosphate homeostasis. The Journal of biological chemistry 288:22670–22680.

Brady, S.M., Song, S., Dhugga, K.S., Rafalski, J.A. and Benfey, P.N. (2006). Combining Expression and Comparative Evolutionary Analysis. The COBRA Gene Family. PLANT PHYSIOLOGY 143:172–187.

Briat, J.-F., Rouached, H., Tissot, N., Gaymard, F. and Dubos, C. (2015). Integration of P, S, Fe, and Zn nutrition signals in Arabidopsis thaliana: potential involvement of PHOSPHATE STARVATION RESPONSE 1 (PHR1). Frontiers in Plant Science 6:290–16.

Castaings, L., Camargo, A., Pocholle, D., Gaudon, V., Texier, Y., Boutet-Mercey, S., Taconnat, L., et al. (2009). The nodule inception-like protein 7 modulates nitrate sensing and metabolism in Arabidopsis. The Plant Journal 57:426–435.

Chevalier, F., Pata, M., Nacry, P., Doumas, P. and Rossignol, M. (2003). Effects of phosphate availability on the root system architecture: large-scale analysis of the natural variation between Arabidopsis accessions. Plant, Cell & Environment 26:1839–1850.

Chiou, T.-J. and Lin, S.-I. (2011). Signaling network in sensing phosphate availability in plants. Annual review of plant biology 62:185–206.

Conte, S.S. and Walker, E.L. (2011). Transporters contributing to iron trafficking in plants. Molecular Plant 4:464–476.

Crawford, N.M. and Glass, A.D.M. (1998). Molecular and physiological aspects of nitrate uptake in plants. Trends in plant science 3:389–395.

De Smet, R. and Marchal, K. (2010). Advantages and limitations of current network inference methods. Nature reviews. Microbiology 8:717–729.

Duan, K., Yi, K., Dang, L., Huang, H., Wu, W. and Wu, P. (2008). Characterization of a sub-family of Arabidopsis genes with the SPX domain reveals their diverse functions in plant tolerance to phosphorus starvation. The Plant Journal 54:965–975.

Fan, W. and Dong, X. (2002). In vivo interaction between NPR1 and transcription factor TGA2 leads to salicylic acid-mediated gene activation in Arabidopsis. The Plant Cell 14:1377–1389.

Food and Agriculture Organization Statistics. Available at: http://faostat3.fao.org/download/Q/QC/E.

Forde, B.G. (2014). Nitrogen signalling pathways shaping root system architecture: an update. Current Opinion in Plant Biology 21:30–36.

Gatz, C. (2013). From Pioneers to Team Players: TGA Transcription Factors Provide a Molecular Link Between Different Stress Pathways. Molecular plant-microbe interactions : MPMI 26:151–159.

Giehl, R.F.H. and Wirén, von, N. (2014). Root nutrient foraging. PLANT PHYSIOLOGY 166:509–517.

Giehl, R.F.H., Gruber, B.D. and Wirén, von, N. (2013). It’s time to make changes: modulation of root system architecture by nutrient signals. Journal of Experimental Botany 65:769–778.

Gruber, B.D., Giehl, R.F.H., Friedel, S. and Wirén, von, N. (2013). Plasticity of the Arabidopsis root system under nutrient deficiencies. PLANT PHYSIOLOGY 163:161–179.

Gu, M., Chen, A., Sun, S. and Xu, G. (2016). Complex Regulation of Plant Phosphate Transporters and the Gap between Molecular Mechanisms and Practical Application: What Is Missing? Molecular Plant 9:396–416.

Guan, P. (2017). Dancing with Hormones: A Current Perspective of Nitrate Signaling and Regulation in Arabidopsis. Frontiers in Plant Science 8:1697.

Gutiérrez-Alanís, D., Yong-Villalobos, L., Jiménez-Sandoval, P., Alatorre-Cobos, F., Oropeza-Aburto, A., Mora-Macías, J., Sánchez-Rodríguez, F., et al. (2017). Phosphate Starvation-Dependent Iron Mobilization Induces CLE14 Expression to Trigger Root Meristem Differentiation through CLV2/PEPR2 Signaling. Developmental Cell 41:555–570.e3.

Hofmann, N.R. (2012). Nicotianamine in zinc and iron homeostasis. The Plant Cell 24:373–373.

Hulsen, T., de Vlieg, J. and Alkema, W. (2008). BioVenn - a web application for the comparison and visualization of biological lists using area-proportional Venn diagrams. BMC Genomics 9:488.

Huynh-Thu, V.A., Irrthum, A., Wehenkel, L. and Geurts, P. (2010). Inferring regulatory networks from expression data using tree-based methods. Isalan, M. (ed.). PLoS ONE 5:e12776.

International Wheat Genome Sequencing Consortium (IWGSC), IWGSC RefSeq principal investigators:, Keller, B., IWGSC whole-genome assembly principal investigators:, Distelfeld, A., Eversole, K., Whole-genome sequencing and assembly:, et al. (2018). Shifting the limits in wheat research and breeding using a fully annotated reference genome. Science (New York, N.Y.) 361:eaar7191.

Jakoby, M., Weisshaar, B., Dröge-Laser, W., Vicente-Carbajosa, J., Tiedemann, J., Kroj, T. and Parcy, F. (2002). bZIP transcription factors in Arabidopsis. Trends in plant science 7:106–111.

Jones, D.T., Taylor, W.R. and Thornton, J.M. (1992). The rapid generation of mutation data matrices from protein sequences. Bioinformatics (Oxford, England) 8:275–282.

Kant, S., Peng, M. and Rothstein, S.J. (2011). Genetic regulation by NLA and microRNA827 for maintaining nitrate-dependent phosphate homeostasis in arabidopsis. PLoS Genetics 7:e1002021–11.

Kapulnik, Y. and Koltai, H. (2016). Fine-tuning by strigolactones of root response to low phosphate. Journal of Integrative Plant Biology 58:203–212.

Kellermeier, F., Armengaud, P., Seditas, T.J., Danku, J., Salt, D.E. and Amtmann, A. (2014). Analysis of the Root System Architecture of Arabidopsis Provides a Quantitative Readout of Crosstalk between Nutritional Signals. The Plant Cell 26:1480–1496.

Khan, A., Fornes, O., Stigliani, A., Gheorghe, M., Castro-Mondragon, J.A., van der Lee, R., Bessy, A., et al. (2018). JASPAR 2018: update of the open-access database of transcription factor binding profiles and its web framework. Nucleic acids research 46:D1284–D1284.

Kiba, T., Inaba, J., Kudo, T., Ueda, N., Konishi, M., Mitsuda, N., Takiguchi, Y., Kondou, Y., Yoshizumi, T., Ohme-Takagi, M., Matsui, M., Yano, K., Yanagisawa, S. and Sakakibara, H. (2018). Repression of Nitrogen Starvation Responses by Members of the Arabidopsis GARP-Type Transcription Factor NIGT1/HRS1 Subfamily. The Plant Cell 30:925–945.

Kieber, J.J. and Schaller, G.E. (2018). Cytokinin signaling in plant development. Development (Cambridge, England) 145:dev149344.

Koen, E., Besson-Bard, A., Duc, C., Astier, J., Gravot, A., Richaud, P., Lamotte, O., et al. (2013). Arabidopsis thaliana nicotianamine synthase 4 is required for proper response to iron deficiency and to cadmium exposure. Plant science : an international journal of experimental plant biology 209:1–11.

Krapp, A., David, L.C., Chardin, C., Girin, T., Marmagne, A., Leprince, A.-S., Chaillou, S., et al. (2014). Nitrate transport and signalling in Arabidopsis. Journal of Experimental Botany 65:789–798.

Krouk, G. (2017). Hormones and nitrate: a two-way connection. Plant Molecular Biology 91:599–606.

Krouk, G., Mirowski, P., LeCun, Y., Shasha, D.E. and Coruzzi, G.M. (2010). Predictive network modeling of the high-resolution dynamic plant transcriptome in response to nitrate. Genome biology 11:R123.

Kumar, R.K., Chu, H.-H., Abundis, C., Vasques, K., Rodriguez, D.C., Chia, J.-C., Huang, R., et al. (2017). Iron-Nicotianamine Transporters Are Required for Proper Long Distance Iron Signaling. PLANT PHYSIOLOGY 175:1254–1268.

Landrein, B., Formosa-Jordan, P., Malivert, A., Schuster, C., Melnyk, C.W., Yang, W., Turnbull, C., et al. (2018). Nitrate modulates stem cell dynamics in Arabidopsis shoot meristems through cytokinins. Proceedings of the National Academy of Sciences of the United States of America 115:1382–1387.

Langfelder, P. and Horvath, S. (2008). WGCNA: an R package for weighted correlation network analysis. BMC bioinformatics 9:559.

Lavenus, J., Goh, T., Roberts, I., Guyomarc’h, S., Lucas, M., De Smet, I., Fukaki, H., et al. (2013). Lateral root development in Arabidopsis: fifty shades of auxin. Trends in plant science 18:450–458.

Lebel, E., Heifetz, P., Thorne, L., Uknes, S., Ryals, J. and Ward, E. (1998). Functional analysis of regulatory sequences controlling PR-1 gene expression in Arabidopsis. The Plant Journal 16:223–233.

Lee, I., Ambaru, B., Thakkar, P., Marcotte, E.M. and Rhee, S.Y. (2010). Rational association of genes with traits using a genome-scale gene network for Arabidopsis thaliana. Nature biotechnology 28:149–156.

Li, B. and Dewey, C.N. (2011). RSEM: accurate transcript quantification from RNA-Seq data with or without a reference genome. BMC bioinformatics 12:323.

Li, W.-F., Perry, P.J., Prafulla, N.N. and Schmidt, W. (2010). Ubiquitin-specific protease 14 (UBP14) is involved in root responses to phosphate deficiency in Arabidopsis. Molecular Plant 3:212–223.

Liu, H., Yang, H., Wu, C., Feng, J., Liu, X., Qin, H. and Wang, D. (2009). Overexpressing HRS1 confers hypersensitivity to low phosphate-elicited inhibition of primary root growth in Arabidopsis thaliana. Journal of Integrative Plant Biology 51:382–392.

Liu, J., Yang, L., Luan, M., Wang, Y., Zhang, C., Zhang, B., Shi, J., et al. (2015). A vacuolar phosphate transporter essential for phosphate homeostasis in Arabidopsis. Proceedings of the National Academy of Sciences of the United States of America 112:E6571–8.

Liu, K.-H., Niu, Y., Konishi, M., Wu, Y., Du, H., Chung, H.S., Li, L., et al. (2017). Discovery of nitrate–CPK–NLP signalling in central nutrient–growth networks. Nature Publishing Group 545:311–316.

Love, M.I., Huber, W. and Anders, S. (2014). Moderated estimation of fold change and dispersion for RNA-seq data with DESeq2. Genome biology 15:550.

Maeda, Y., Konishi, M., Kiba, T., Sakuraba, Y., Sawaki, N., Kurai, T., Ueda, Y., et al. (2018). A NIGT1-centred transcriptional cascade regulates nitrate signalling and incorporates phosphorus starvation signals in Arabidopsis. Nature Communications:1–14.

Maere, S., Heymans, K. and Kuiper, M. (2005). BiNGO: a Cytoscape plugin to assess overrepresentation of gene ontology categories in biological networks. Bioinformatics (Oxford, England) 21:3448–3449.

Marchive, C., Roudier, F., Castaings, L., Bréhaut, V., Blondet, E., Colot, V., Meyer, C., et al. (2013). Nuclear retention of the transcription factor NLP7 orchestrates the early response to nitrate in plants. Nature Communications 4:1713–9.

Marcussen, T., Sandve, S.R., Heier, L., Spannagl, M., Pfeifer, M., International Wheat Genome Sequencing Consortium,, Jakobsen, K.S., et al. (2014). Ancient hybridizations among the ancestral genomes of bread wheat. Science (New York, N.Y.) 345:1250092–1250092.

Marschner, H. (2011). Marschner’s Mineral Nutrition of Higher Plants. 3rd ed. Academic Press.

McClure, B.A., Hagen, G., Brown, C.S., Gee, M.A. and Guilfoyle, T.J. (1989). Transcription, organization, and sequence of an auxin-regulated gene cluster in soybean. The Plant Cell 1:229–239.

Medici, A., Marshall-Colon, A., Ronzier, E., Szponarski, W., Wang, R., Gojon, A., Crawford, N.M., et al. (2015). AtNIGT1/HRS1 integrates nitrate and phosphate signals at the Arabidopsis root tip. Nature Communications 6:1–11.

Mora-Macías, J., Ojeda-Rivera, J.O., Gutiérrez-Alanís, D., Yong-Villalobos, L., Oropeza-Aburto, A., Raya-González, J., Jiménez-Domínguez, G., et al. (2017). Malate-dependent Fe accumulation is a critical checkpoint in the root developmental response to low phosphate. Proceedings of the National Academy of Sciences 114:E3563–E3572.

Nilsson, L., Müller, R. and Nielsen, T.H. (2007). Increased expression of the MYB-related transcription factor, PHR1, leads to enhanced phosphate uptake in Arabidopsis thaliana. Plant, Cell & Environment 30:1499–1512.

Niu, Y.F., Chai, R.S., Jin, G.L., Wang, H., Tang, C.X. and Zhang, Y.S. (2013). Responses of root architecture development to low phosphorus availability: a review. Annals of botany 112:391–408.

Obayashi, T., Aoki, Y., Tadaka, S., Kagaya, Y. and Kinoshita, K. (2018). ATTED-II in 2018: A Plant Coexpression Database Based on Investigation of the Statistical Property of the Mutual Rank Index. Plant and Cell Physiology 59:440–440.

Pal, S., Kisko, M., Dubos, C., Lacombe, B., Berthomieu, P., Krouk, G. and Rouached, H. (2017). TransDetect identifies a new regulatory module controlling phosphate accumulation. PLANT PHYSIOLOGY:pp.00568.2017–11.

Patro, R., Duggal, G., Love, M.I., Irizarry, R.A. and Kingsford, C. (2017). Salmon provides fast and bias-aware quantification of transcript expression. Nature methods 14:417–419.

Péret, B., Desnos, T., Jost, R., Kanno, S., Berkowitz, O. and Nussaume, L. (2014). Root architecture responses: in search of phosphate. PLANT PHYSIOLOGY 166:1713–1723.

Pfeifer, M., Kugler, K.G., Sandve, S.R., Zhan, B., Rudi, H., Hvidsten, T.R., International Wheat Genome Sequencing Consortium,, et al. (2014). Genome interplay in the grain transcriptome of hexaploid bread wheat. Science (New York, N.Y.) 345:1250091–1250091.

Piñeros, M.A., Cançado, G.M.A. and Kochian, L.V. (2008). Novel properties of the wheat aluminum tolerance organic acid transporter (TaALMT1) revealed by electrophysiological characterization in Xenopus Oocytes: functional and structural implications. PLANT PHYSIOLOGY 147:2131–2146.

Powell, J.J., Fitzgerald, T.L., Stiller, J., Berkman, P.J., Gardiner, D.M., Manners, J.M., Henry, R.J., et al. (2017). The defence-associated transcriptome of hexaploid wheat displays homoeolog expression and induction bias. Plant biotechnology journal 15:533–543.

Puga, M.I., Mateos, I., Charukesi, R., Wang, Z., Franco-Zorrilla, J.M., de Lorenzo, L., Irigoyen, M.L., et al. (2014). SPX1 is a phosphate-dependent inhibitor of PHOSPHATE STARVATION RESPONSE 1 in Arabidopsis. Proceedings of the National Academy of Sciences 111:14947–14952.

Roudier, F. (2002). The COBRA Family of Putative GPI-Anchored Proteins in Arabidopsis. A New Fellowship in Expansion. PLANT PHYSIOLOGY 130:538–548.

Rubin, G., Tohge, T., Matsuda, F., Saito, K. and Scheible, W.-R. (2009). Members of the LBD family of transcription factors repress anthocyanin synthesis and affect additional nitrogen responses in Arabidopsis. The Plant Cell 21:3567–3584.

Ruffel, S., Krouk, G., Ristova, D., Shasha, D., Birnbaum, K.D. and Coruzzi, G.M. (2011). Nitrogen economics of root foraging: transitive closure of the nitrate-cytokinin relay and distinct systemic signaling for N supply vs. demand. Proceedings of the National Academy of Sciences of the United States of America 108:18524–18529.

Ruffel, S., Poitout, A., Krouk, G., Coruzzi, G.M. and Lacombe, B. (2016). Long-distance nitrate signaling displays cytokinin dependent and independent branches. Journal of Integrative Plant Biology 58:226–229.

Sakakibara, H., Takei, K. and Hirose, N. (2006). Interactions between nitrogen and cytokinin in the regulation of metabolism and development. Trends in plant science 11:440–448.

Sawaki, N., Tsujimoto, R., Shigyo, M., Konishi, M., Toki, S., Fujiwara, T. and Yanagisawa, S. (2013). A Nitrate-Inducible GARP Family Gene Encodes an Auto-Repressible Transcriptional Repressor in Rice. Plant and Cell Physiology 54:506–517.

Schachtman, D., Reid, R. and Ayling, S. (1998). Phosphorus Uptake by Plants: From Soil to Cell. PLANT PHYSIOLOGY 116:447–453.

Scheible, W.-R., Morcuende, R., Czechowski, T., Fritz, C., Osuna, D., Palacios-Rojas, N., Schindelasch, D., et al. (2004). Genome-wide reprogramming of primary and secondary metabolism, protein synthesis, cellular growth processes, and the regulatory infrastructure of Arabidopsis in response to nitrogen. PLANT PHYSIOLOGY 136:2483–2499.

Schindelman, G., Morikami, A., Jung, J., Baskin, T.I., Carpita, N.C., Derbyshire, P., McCann, M.C. and Benfey, P.N. (2001). COBRA encodes a putative GPI-anchored protein, which is polarly localized and necessary for oriented cell expansion in Arabidopsis. :1–14.

Schuler, M., Rellán-Álvarez, R., Fink-Straube, C., Abadía, J. and Bauer, P. (2012). Nicotianamine Functions in the Phloem-Based Transport of Iron to Sink Organs, in Pollen Development and Pollen Tube Growth in Arabidopsis. The Plant Cell 24:2380–2400.

Secco, D., Jabnoune, M., Walker, H., Shou, H., Wu, P., Poirier, Y. and Whelan, J. (2013). Spatio-Temporal Transcript Profiling of Rice Roots and Shoots in Response to Phosphate Starvation and Recovery. The Plant Cell 25:4285–4304.

Shahzad, Z. and Amtmann, A. (2017). Food for thought: how nutrients regulate root system architecture. Current Opinion in Plant Biology 39:80–87.

Shahzad, Z., Kellermeier, F., Armstrong, E.M., Rogers, S., Lobet, G., Amtmann, A. and Hills, A. (2018). EZ-Root-VIS: A Software Pipeline for the Rapid Analysis and Visual Reconstruction of Root System Architecture. PLANT PHYSIOLOGY 177:1368–1381.

Shannon, P., Markiel, A., Ozier, O., Baliga, N.S., Wang, J.T., Ramage, D., Amin, N., et al. (2003). Cytoscape: a software environment for integrated models of biomolecular interaction networks. Genome Research 13:2498–2504.

Li Shen and Mount Sinai (2013). GeneOverlap: Test and visualize gene overlaps. R package version 1.8.0. http://shenlab-sinai.github.io/shenlab-sinai/

Shukla, D., Rinehart, C.A. and Sahi, S.V. (2017). Comprehensive study of excess phosphate response reveals ethylene mediated signaling that negatively regulates plant growth and development. Nature Publishing Group 7:3074.

Smith, S. and De Smet, I. (2012). Root system architecture: insights from Arabidopsis and cereal crops. Philosophical transactions of the Royal Society of London. Series B, Biological sciences 367:1441–1452.

Subramaniam, R., Desveaux, D., Spickler, C., Michnick, S.W. and Brisson, N. (2001). Direct visualization of protein interactions in plant cells. Nature biotechnology 19:769–772.

Sun, H., Tao, J., Liu, S., Huang, S., Chen, S., Xie, X., Yoneyama, K., et al. (2014). Strigolactones are involved in phosphate- and nitrate-deficiency-induced root development and auxin transport in rice. Journal of Experimental Botany 65:6735–6746.

Takahashi, M., Terada, Y., Nakai, I., Nakanishi, H., Yoshimura, E., Mori, S. and Nishizawa, N.K. (2003). Role of Nicotianamine in the Intracellular Delivery of Metals and Plant Reproductive Development. Plant Cell 15:1263.

Tamura, K., Stecher, G., Peterson, D., Filipski, A. and Kumar, S. (2013). MEGA6: Molecular Evolutionary Genetics Analysis version 6.0. Molecular biology and evolution 30:2725–2729.

Thompson, J.D., Higgins, D.G. and Gibson, T.J. (1994). CLUSTAL W: improving the sensitivity of progressive multiple sequence alignment through sequence weighting, position-specific gap penalties and weight matrix choice. Nucleic acids research 22:4673–4680.

Tian, H., De Smet, I. and Ding, Z. (2014). Shaping a root system: regulating lateral versus primary root growth. Trends in plant science 19:426–431.

Tilman, D., Cassman, K.G., Matson, P.A., Naylor, R. and Polasky, S. (2002). Agricultural sustainability and intensive production practices. Nature 418:671–677.

Usadel, B., Obayashi, T., Mutwil, M., Giorgi, F.M., Bassel, G.W., Tanimoto, M., Chow, A., et al. (2009). Co-expression tools for plant biology: opportunities for hypothesis generation and caveats. Plant, Cell & Environment 32:1633–1651.

Vance, C.P. (2001). Symbiotic nitrogen fixation and phosphorus acquisition. Plant nutrition in a world of declining renewable resources. PLANT PHYSIOLOGY 127:390–397.

Wang, C., Yue, W., Ying, Y., Wang, S., Secco, D., Liu, Y., Whelan, J., et al. (2015). Rice SPX-Major Facility Superfamily3, a Vacuolar Phosphate Efflux Transporter, Is Involved in Maintaining Phosphate Homeostasis in Rice. PLANT PHYSIOLOGY 169:2822–2831.

Wang, R., Guegler, K., LaBrie, S.T. and Crawford, N.M. (2000). Genomic analysis of a nutrient response in Arabidopsis reveals diverse expression patterns and novel metabolic and potential regulatory genes induced by nitrate. The Plant Cell 12:1491–1509.

Wang, R., Okamoto, M., Xing, X. and Crawford, N.M. (2003). Microarray analysis of the nitrate response in Arabidopsis roots and shoots reveals over 1,000 rapidly responding genes and new linkages to glucose, trehalose-6-phosphate, iron, and sulfate metabolism. PLANT PHYSIOLOGY 132:556–567.

Waters, B.M., Chu, H.-H., Didonato, R.J., Roberts, L.A., Eisley, R.B., Lahner, B., Salt, D.E., et al. (2006). Mutations in Arabidopsis yellow stripe-like1 and yellow stripe-like3 reveal their roles in metal ion homeostasis and loading of metal ions in seeds. PLANT PHYSIOLOGY 141:1446–1458.

Wiren N, von, Klair, S., Bansal, S., Briat, J., Khodr, H., Shioiri, T., Leigh, R., et al. (1999). Nicotianamine chelates both FeIII and FeII. Implications for metal transport in plants. PLANT PHYSIOLOGY 119:1107–1114.

Wolfe, C.J., Kohane, I.S. and Butte, A.J. (2005). Systematic survey reveals general applicability of ‘guilt-by-association’ within gene coexpression networks. BMC bioinformatics 6:227.

Yu, P., Gutjahr, C., Li, C. and Hochholdinger, F. (2016). Genetic Control of Lateral Root Formation in Cereals. Trends in plant science 21:951–961.

Zhang, W.-H., Ryan, P.R., Sasaki, T., Yamamoto, Y., Sullivan, W. and Tyerman, S.D. (2008). Characterization of the TaALMT1 Protein as an Al3+-Activated Anion Channel in Transformed Tobacco (Nicotiana tabacum L.) Cells. Plant and Cell Physiology 49:1316–1330.

Zhang, X., Davidson, E.A., Mauzerall, D.L., Searchinger, T.D., Dumas, P. and Shen, Y. (2015). Managing nitrogen for sustainable development. Nature 528:51–59.

